# Metabolic cross-talk promotes persistence of *Enterococcus* in a model of polymicrobial catheter-associated urinary tract infection

**DOI:** 10.1101/2025.11.19.689321

**Authors:** Zongsen Zou, Jerome S. Pinkner, Chloe L. P. Obernuefemann, Kent R. Kleinschmidt, Denise A. Sanick, Suzanne M. Hickerson, Karen W. Dodson, Jeffrey P. Henderson, Scott J. Hultgren, Michael G. Caparon

## Abstract

Catheter-associated urinary tract infections (CAUTI) account for 70%∼80% of urinary tract infections (UTI) and can lead to adverse outcomes. Most CAUTIs are polymicrobial with resilient communities maintaining a consistent composition of species over time, despite antibiotic treatment and catheter replacement. However, the mechanisms promoting persistence are poorly understood. Here we examine how a chemical interaction between Gram-positive *Enterococcus faecalis* and Gram-negative *Klebsiella pneumoniae* can explain their high rate of co-occurrence on long-term indwelling urinary catheters. Sequence analyses of longitudinal isolates from several human patients co-infected with *E. faecalis* and *K. pneumoniae* revealed that despite frequent replacement, catheters became re-colonized with the same or a nearly identical consortium of strains throughout the study collection period. Using artificial urine medium (AUM), monoculture revealed that the *K. pneumoniae* isolates grew robustly and formed biofilm, while the *E. faecalis* isolates grew poorly and did not form biofilm. However, co-culture of paired isolates resulted in enhanced *E. faecalis* growth and biofilm, which could be reproduced by supplementing *E. faecalis* with cell-free *K. pneumoniae* conditioned AUM supernatant (KpAUMSup). Analyses using comparative transcriptomics, mutant strains and chemical inhibitors with cell culture and murine CAUTI models revealed that: **i)** KpAUMSup, but not AUM, stimulated expression of the *E. faecalis* Fsr quorum-sensing system; **ii)** Fsr was required for *E. faecalis* to respond to KpAUMSup; **iii)** *E. faecalis* cultured in KpAUMSup was more efficient in initiating CAUTI; and **iv)** Disruption of Fsr inhibited initiation of CAUTI. This interspecies signaling may help explain the high rate of co-colonization of these CAUTI pathogens and highlights new therapeutic strategies to treat polymicrobial CAUTI.

## Introduction

It is well-appreciated that many infectious diseases are caused by a consortium of diverse species interacting within the context of an integrated bacterial community as opposed to a single etiological entity. What is less well-appreciated are the underlying mechanisms that dictate how polymicrobial communities are assembled during infection, how they are resilient despite host defenses and medical treatment, and how interactions between members of the community can influence its pathogenic potential, a concept known as nososymbiocity (*1, 2*).

Ecological studies have indicated that community assembly is typically the result of non-stochastic processes influenced by several positive and negative strategies, which can include **i)** facultative mutualism, where a keystone species modifies an environment to facilitate the proliferation of symbiotic partners (*3, 4*); **ii)** selective elimination of competitor species through the production of targeted antimicrobial agents (*5*); **iii)** trophic interactions where species are interconnected through the exchange of metabolites (*6*); and **iv)** interbacterial communication, where the production and sensing of dedicated signaling molecules coordinates community behavior (*7, 8*). Understanding how any of these strategies can contribute to the pathogenesis of polymicrobial infections may lead to novel therapeutic strategies for their treatment and prevention.

CAUTIs are the most common cause of hospital-acquired infections worldwide and account for up to 1 million cases in the USA alone and up to $4.5 billion in additional health care costs. As many as 15-25% of all hospitalized patients receive a urinary catheter and the risk of bacterial colonization of the catheter increases 3-7% per day following the implantation with colonization rates approaching 100% in long-term catheterized patients (*9–12*). While CAUTI can be caused by a diverse array of Gram-positive and Gram-negative pathogens, the majority are polymicrobial, which promotes antibiotic resistance, complicates treatment, and often results in increased mortality rate (*13–15*). Thus, an understanding of the mechanisms that promote polymicrobial CAUTI leading to enhanced nososymbiocity, will be critical for the development of novel strategies to prevent polymicrobial CAUTI in long-term catheterized patients to produce favorable clinical outcomes. Our recent examination of catheter and urine samples recovered from long-term catheterized human patients revealed that the formation of resilient polymicrobial biofilm communities (typically composed of between 2-3 bacterial species) was often the result of a non-stochastic process, as we identified numerous positive and negative genus-level co-associations that occurred at frequencies that were either more or less than expected by chance (*16*). These associations were resilient, where despite antibiotic treatment and frequent catheter replacement, the composition of microbial species in polymicrobial biofilm within a particular catheterized individual remained constant across multiple ∼monthly collection periods. Overall, the most common species identified in our clinical study was the Gram-positive coccus *E. faecalis*, which was positively co-associated with one of several Gram-negative enteric species, such as *E. coli* and *K. pneumoniae* (*16*). In-depth examination of the co-association of paired longitudinal isolates of *E. faecalis* and *E. coli* revealed an apparent functional association through their spatial co-localization on recovered human catheters and from *in vitro* co-culture studies using an artificial urine medium (AUM) where *E. faecalis* achieved enhanced growth in the presence of *E. coli*.

In the present study, we examine functional mechanisms that can promote polymicrobial CAUTI through analysis of the interaction between *E. faecalis* and *K. pneumoniae*, which our prior study indicated to have both a significantly high co-occurrence rate and a stable frequency of longitudinal co-colonization in long-term catheterized patients. Our studies revealed that a cell-free soluble factor(s) generated by *K. pneumoniae* during growth in AUM can stimulate the Fsr quorum-sensing system of *E. faecalis* to enhance its ability to grow and form biofilm *in vitro* and enhance its efficiency in initiating infection in both a cell culture model and in a murine model of CAUTI. These findings lay the groundwork for the development of novel therapeutic strategies that can perturb the formation and stability of problematic polymicrobial communities to treat and prevent CAUTI.

## Results

### Persistent polymicrobial CAUTI caused by the same lineages of *E. faecalis* and *K. pneumoniae*

To examine the interaction between *E. faecalis* and *K. pneumoniae*, we chose a collection of longitudinal isolates from two human patients that our prior study indicated were persistently co-colonized by these two organisms (*16*). A limitation of our prior study is that phylogenies based on 16S sequencing lacked the resolution to discriminate between lineages at the sub-species level. Therefore, to characterize community stability at the sub-species level, 9 and 11 longitudinal pairs of isolates from Patients 82 and 85, respectively, were subjected to whole genome sequencing. Unsupervised hierarchical phylogenetic clustering (Fig. 1, A and B) and genomic comparison using average nucleotide identity (ANI) (*E. faecalis*, ANI ≥ 99.93%; *K. pneumoniae* ANI ≥ 99.98%; tables S1 and S2) revealed that both the *E. faecalis* and *K. pneumoniae* isolates from Patient 85 were genetically closely related to each other. Additionally, antibiotic resistant genes analysis revealed that the *K. pneumoniae* isolates (fig. S1B, table S7), but not the *E. faecalis* isolates (fig. S1A, table S6), were identified with stable antibiotic resistance profiles over time. For Patient 82, all *K. pneumoniae* isolates were observed with genetic closeness by phylogenetic (Fig. 1C) and ANI (ANI ≥ 99.98%, table S5) analyses, and had an identical antibiotic resistance profile (fig. S1C, table S9). In contrast, phylogenetic clustering of *E. faecalis* isolates from Patient 82 separated into two clades (Fig. 1C), unlike the other three groups of isolates. These two clades were each highly homogeneous (ANI ≥ 99.97%, ANI ≥ 99.99%; tables S3 and S4) and were identified with the same antibiotic resistance profiles (fig. S1C, table S8). Based on limitations associated with sample collection in our prior study (only a single colony for *E. faecalis* was characterized at each time point) (*16*), it is not possible to determine if the two lineages stably co-existed over time or were in a dynamic mutually exclusive relationship. However, this analysis demonstrates that despite the frequent removal and replacement of catheters, these two patients each remained colonized by a polymicrobial community consisting of the same or nearly identical genomic lineages of *E. faecalis* and *K. pneumoniae* over an extended period.

**Fig. 1.**
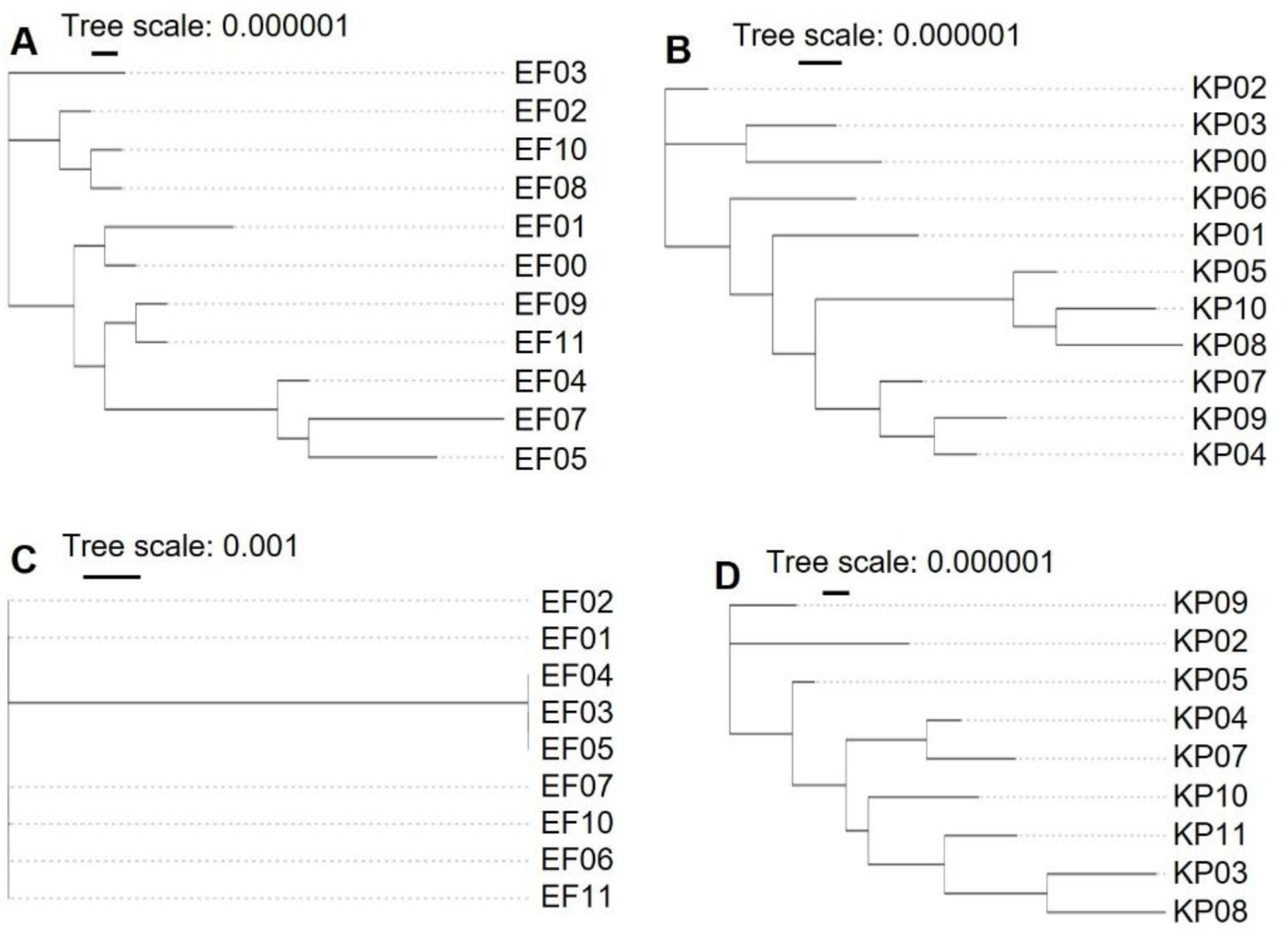
*E. faecalis* and *K. pneumoniae* strains cause recurrent CAUTI in long-term catheterized patients. Phylogenetic distributions of 11 *E. faecalis* (**A**) and 11 *K. pneumoniae* (**B**) clinical isolates from Patient 85, and 9 *E. faecalis* (**C**) and 9 *K. pneumoniae* (**D**) clinical isolates from Patient 82.

### Enhanced *E. faecalis* growth and biofilm formation in co-culture with *K. pneumoniae*

To assess whether interactions that affect growth may contribute to the high frequency of co-colonization, both reference *E. faecalis* (OG1RF) and *K. pneumoniae* (TOP52) strains and individual paired patient isolates were tested in co-culture in several media. In nutrient abundant brain heart infusion (BHI) medium, growth of all strains was robust in monoculture and planktonic yields of each isolate in paired co-culture reached the levels of each isolate in monoculture (figS2, A and B). However, in nutrient-limited AUM, which more closely mimics the growth conditions of catheter isolates in the bladder environment (*17*), all *K. pneumoniae* isolates grew to high densities in both mono- (∼10^7.86^ CFU/mL) and co-cultures (∼10^7.90^ CFU/mL) (Fig. 2B), all *E. faecalis* isolates grew poorly in AUM monoculture (∼10^4.46^ CFU/mL), achieving yields that were up to ∼3 logs lower than in BHI (∼10^7.51^ CFU/mL) (Fig. 2A). These poor planktonic yields were overcome by co-culture with *K. pneumoniae* which enhanced *E. faecalis* growth by ∼2.5 logs (∼10^6.82^ CFU/mL) (Fig. 2A). An identical pattern was observed for co-culture in the presence of a silicone catheter in AUM, where both planktonic (∼10^7.01^ CFU/mL) (Fig. 2C) and sessile (∼10^7.02^ CFU/mL) (Fig. 2D) CFUs of *E. faecalis* OG1RF were increased in co-culture with *K. pneumoniae* TOP52 as compared to monoculture (planktonic = ∼10^3.70^ CFU/mL, sessile = ∼10^3.70^ CFU/mL), while all *K. pneumoniae* TOP52 growth yields remained constant between mono- and co-culture (Fig. 2, E and F). When examined by scanning electron microscopy (SEM), *E. faecalis* in co-culture demonstrated increased chain-lengths and microcolony formation on the catheter surface as compared to mono-culture (Fig. 2, G and H). Immunofluorescent staining of catheters recovered from Patients 85 and 82 also revealed co-localization of *E. faecalis* and *K. pneumoniae* on the catheter surface (Fig. 2, I and J; figs. S3 and S4).

**Fig. 2.**
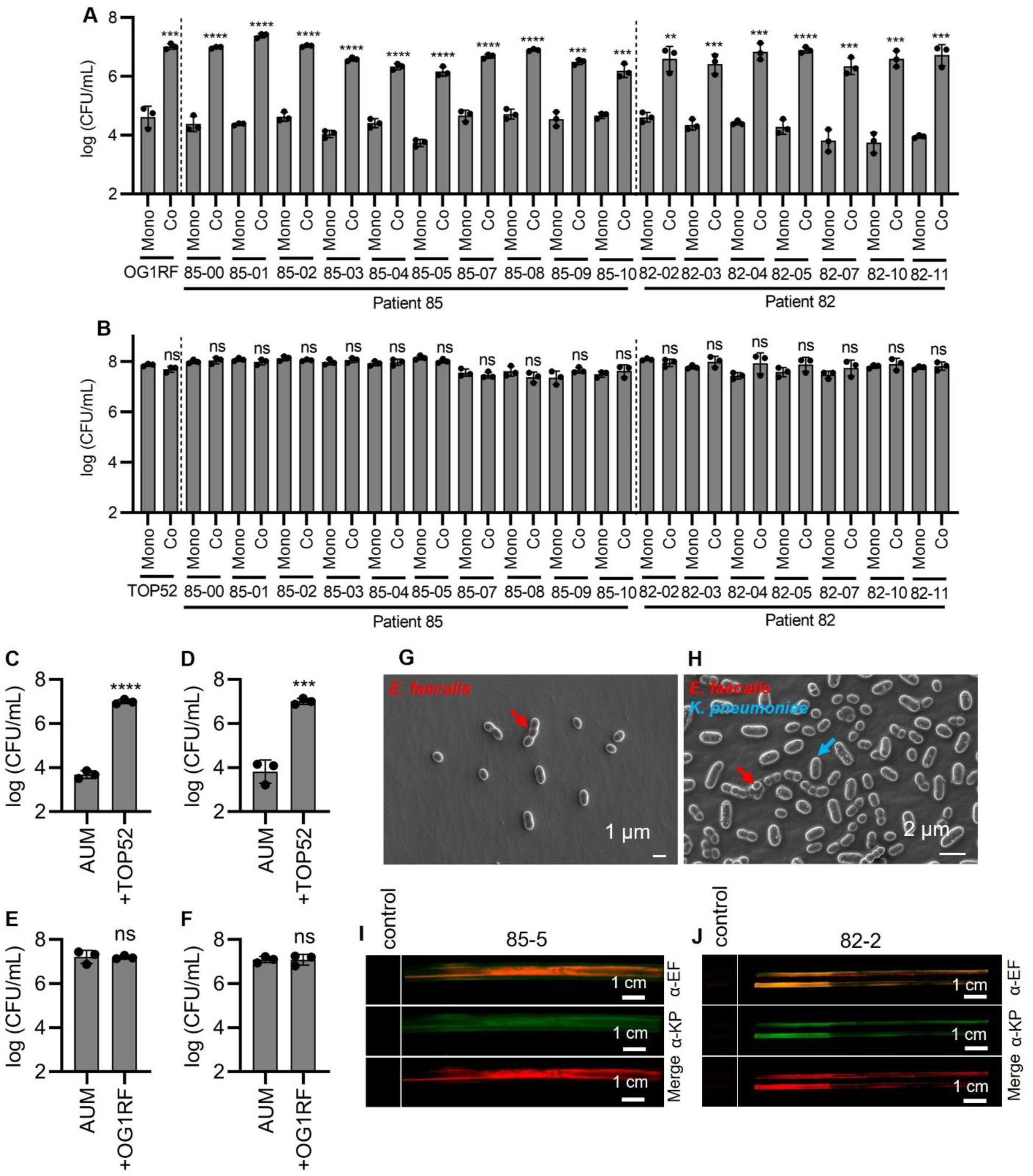
*E. faecalis* achieves enhanced growth and biofilm formation in beneficial interaction with *K. pneumoniae*. (**A** and **B**) Growth of clinical and prototypical *E. faecalis* (**A**) and *K. pneumoniae* (**B**) strains in AUM cultures in microplate, including monoculture of each bacteria (Mono) and co-culture of both bacteria (Co). (**C** and **D**) Growth of planktonic cells in medium (**C**) and sessile cells on silicone catheter surface (**D**) for *E. faecalis* OG1RF cultured in monoculture (AUM) and co-culture with *K. pneumoniae* TOP52 (+TOP52) using AUM. (**E** and **F**) Growth of planktonic cells in medium (**E**) and sessile cells on silicone catheter surface (**F**) for *K. pneumoniae* TOP52 cultured in monoculture (AUM) and co-culture with *E. faecalis* OG1RF (+OG1RF) using AUM. (**G** and **H**) SEM characterization of biofilm formation on catheter surface by *E. faecalis* OG1RF grown in monoculture (**G**) and co-culture with *K. pneumoniae* TOP52 (**H**) using AUM. (**I** and **J**) Immunohistochemistry imaging of *E. faecalis* and *K. pneumoniae* clinical isolates on formalin-fixed catheter portions obtained from the fifth collection period from Patient 85 (**I**) and the second collection period from 82 (**J**). Data represent the mean ± SD derived from three independent experiments. Statistics were performed using a two-tailed unpaired t test with P ≤ 0.05 considered as statistically significant. *P ≤ 0.05, **P < 0.01, ***P < 0.001, ****P < 0.0001, ns indicates not significant.

### *K. pneumoniae* conditioned medium promotes *E. faecalis* growth and biofilm formation

To further investigate the mechanism of interaction between *K. pneumonia* and *E. faecalis*, clinical and reference *E. faecalis* strains were cultured in AUM alone or AUM supplemented with either clinical or reference *K. pneumoniae* strains-conditioned cell-free AUM supernatant (KpAUMSup). The results showed that all strains of *E. faecalis* grown in KpAUMSup achieved significantly enhanced planktonic growth (∼10^7.15^ CFU/mL) to levels obtained in active co-culture with *K. pneumoniae* (Fig. 3A), as relative to growth in AUM alone (∼10^4.66^ CFU/mL). Similarly, *E. faecalis* biofilm formation (A595 = ∼0.85) was enhanced in KpAUMSup (Fig. 3B) as compared to AUM alone (A595 = ∼0.23). Furthermore, KpAUMSup retained its ability to significantly promote *E. faecalis* planktonic growth and biofilm when diluted up to 10^9^- and 10^7^-fold, respectively (fig. S5, A and B). In the silicone catheter assay, KpAUMSup supplementation promoted enhanced planktonic (∼2.52 logs increase) and sessile (∼2.31 logs increase) growth of *E. faecalis* OG1RF as determined by both CFUs (Fig. 3, C and D) and SEM characterization (Fig. 3, E and F). Quantitation of biofilm development in a culture dish assay using immunofluorescent confocal microscopy revealed that KpAUMSup enhanced *E. faecalis* OG1RF biofilm formation in AUM with an ∼6-fold increase of biofilm volume (Fig. 3, G, H, and I). Analysis by filtration indicated that the active factor(s) in KpAUMSup were smaller than 5 kDa (fig. S6) and fractionation by a reverse-phase Amberlite resin and a reverse-phase Phenyl-Hexyl high-performance liquid chromatography (HPLC) column produced two fractions (Fraction 6 and 7) (fig. S7), which significantly enhanced the *E. faecalis* phenotype (fig. S8). Further, the activity of both KpAUMSup and Fractions 6 and 7 was significantly decreased after exposure to 95°C for 24 hrs (fig. S9). Together, these results show that during culture in AUM, *K. pneumoniae* generates a heat-labile small molecular weight extracellular factor(s) that can stimulate *E. faecalis* planktonic growth and biofilm in nutrient-limited media similar to the urine conditions encountered in the bladder.

**Fig. 3.**
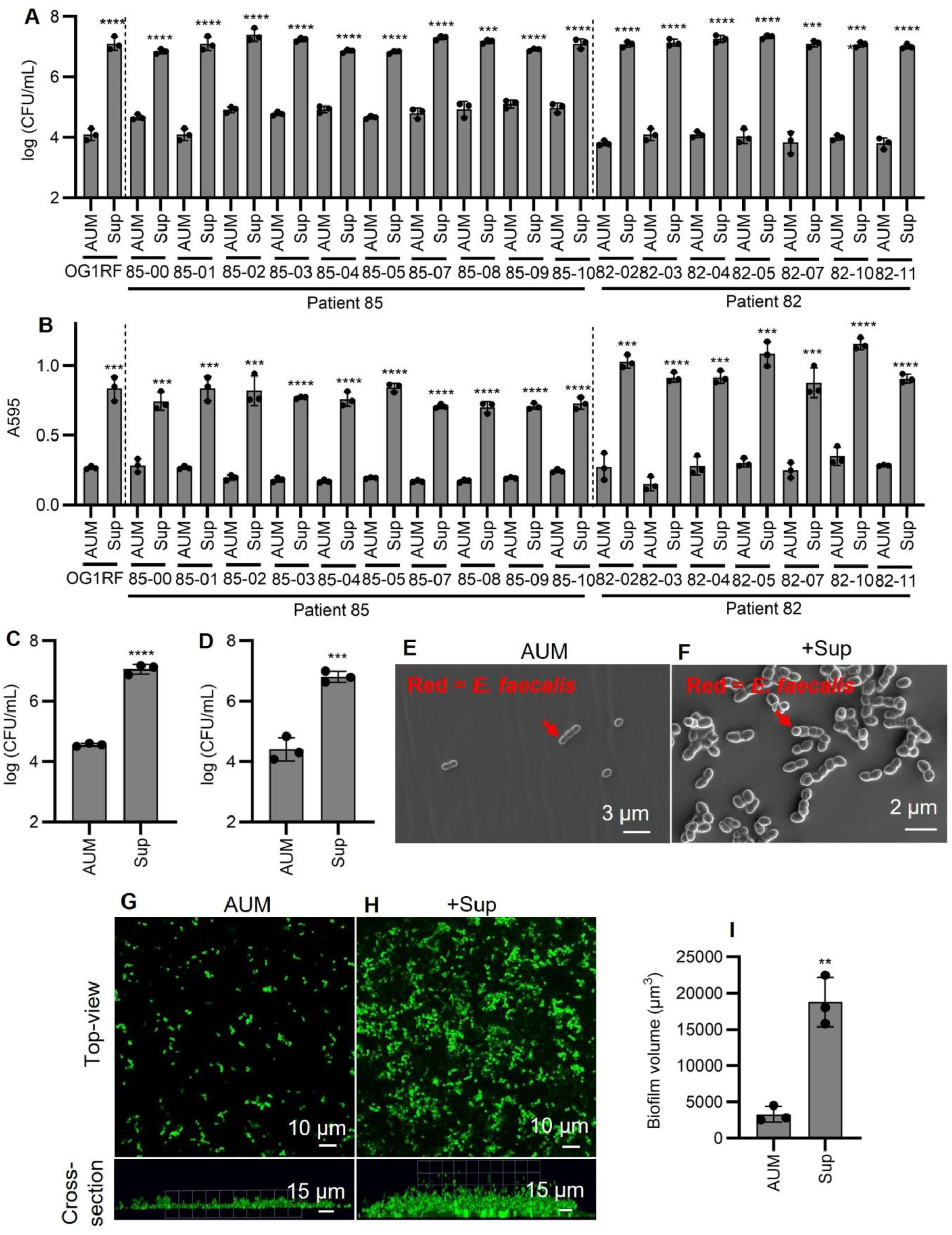
*K. pneumoniae*-generated factors promote *E. faecalis* growth and biofilm formation. (**A** and **B**) Bacterial growth (**A**) and biofilm formation (**B**) of clinical and prototypical *E. faecalis* strains cultured in AUM (AUM) and AUM supplemented with *K. pneumoniae* AUM supernatant (Sup) in microplate. (**C** and **D**) Growth of planktonic cells in medium (**C**) and sessile cells on silicone catheter surface (**D**) for *E. faecalis* OG1RF cultured in AUM (AUM) and AUM supplemented with *K. pneumoniae* TOP52 AUM supernatant (Sup). (**E** and **F**) SEM characterization of biofilm formation on catheter surface by *E. faecalis* OG1RF grown in AUM (AUM) (**E**) and AUM supplemented with *K. pneumoniae* TOP52 AUM supernatant (Sup) (**F**). (**G** to **I**) Immunofluorescent confocal microscopy characterization of *E. faecalis* OG1RF biofilm grown in AUM (AUM) (**G**) and AUM supplemented with *K. pneumoniae* TOP52 AUM supernatant (Sup) (**H**) in peri dishes, followed by biofilm volume quantification using Volocity software (**I**). Data represent the mean ± SD derived from three independent experiments. Statistics were performed using a two-tailed unpaired t test with P ≤ 0.05 considered as statistically significant. *P ≤ 0.05, **P < 0.01, ***P < 0.001, ****P < 0.0001, ns indicates not significant.

### Up-regulation of the *E. faecalis* Fsr quorum sensing system

To decipher the mechanism by which KpAUMSup stimulates both planktonic growth and biofilm in *E. faecalis*, we employed comparative transcriptomics. Our strategy was to examine the effect of KpAUMSup on the *E. faecalis* transcriptome under different conditions that; i) emphasized the importance of growth-associated genes; and ii) emphasized the importance of biofilm-associated genes. The genes of interest would be those that were associated with both growth and biofilm. For analysis of growth-associated pathways, the kinetics of planktonic growth was characterized over time following exposure to KpAUMSup. At 9 hrs post-inoculation, no stimulation of growth (fig. S10A) or biofilm (fig. S10B) was observed. In contrast, by 12 hours post-inoculation, growth was enhanced (fig. S10A) with no increase in biofilm (fig. S10B), suggesting that genes differentially expressed at the 9 hr time point would be those induced to stimulate planktonic growth. A principal component analysis (PCA) of RNA-Seq comparing the *E. faecalis* OG1RF transcriptome following 9 hrs of incubation in the presence or absence of KpAUMSup demonstrated distinct separation of the supplemented and non-supplemented transcriptomes along the PC1 axis (accounting for 70.2% of total variance) (Fig. 4A), indicating that the two groups have distinct transcriptomic profiles. Stringent criteria (|log_2_(FC)| and -log(P) > 99% confidence intervals upper limits) were applied to identify the most significantly regulated genes associated with growth with 40 genes identified as up-regulated (Fig. 4B, E, and F; table S10) and 14 genes identified as down-regulated (Fig. 4B; table S10).

**Fig. 4.**
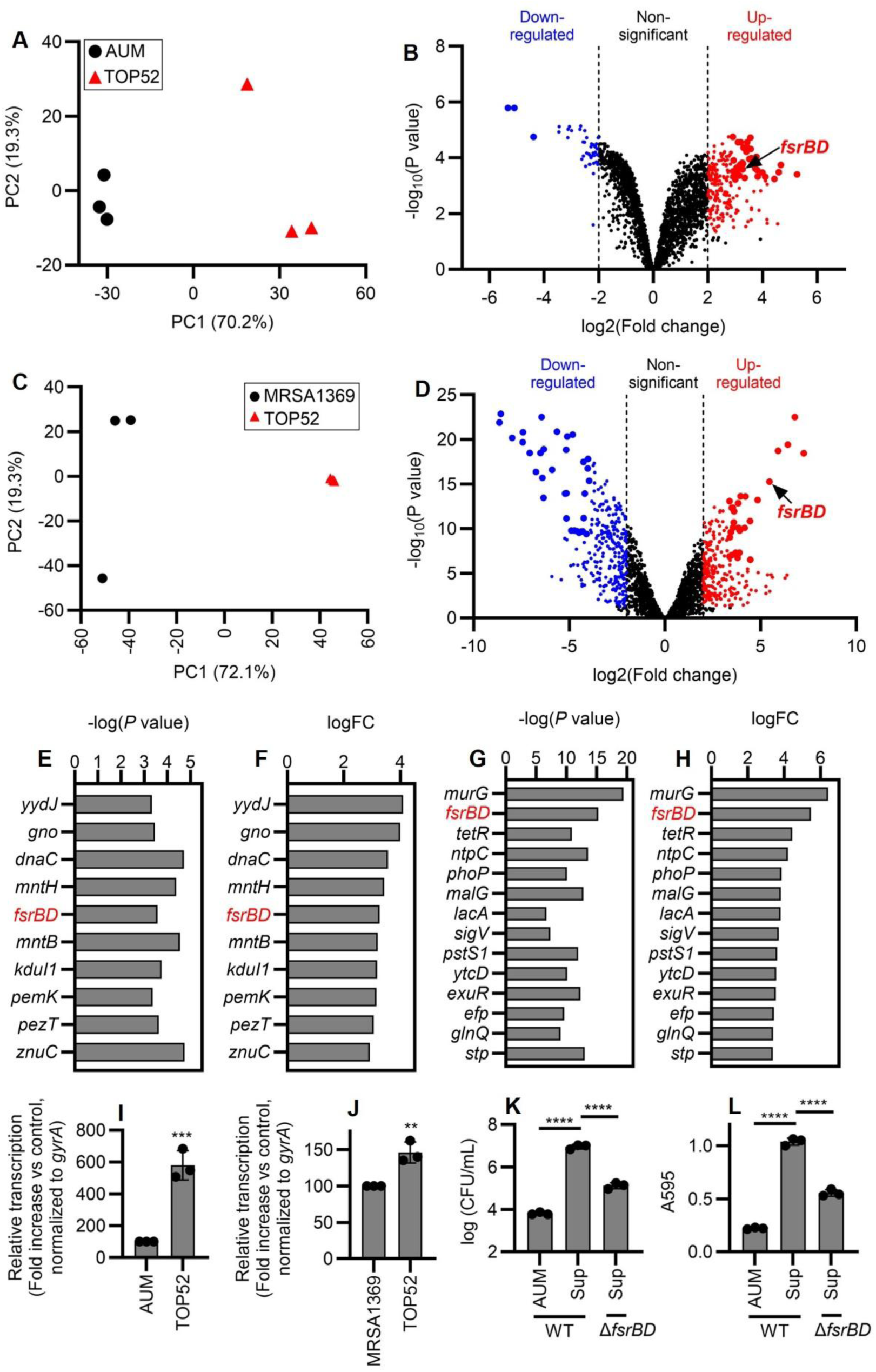
*K. pneumoniae* active factors upregulate *E. faecalis* Fsr quorum sensing system. (**A**) Score plot (PC2∼PC1) of the first (PC1) and second (PC2) principal components in principal component analysis (PCA) showed the clear separation between the two transcriptome groups along the PC1 axis (accounting for 70.2% of total variance between specimens), revealing their different transcriptome profiles. (**B**) A volcano plot identified *E. faecalis* growth-associated differentially expressed genes (DEGs) (|log_2_(FC)| > 2.0 and P < 0.05), with down- and up-regulated genes labelled as blue and red dots, respectively. More stringent criteria (|log2(FC)| and -log(P) > 99% confidence intervals upper limits) were applied to identify the most significantly regulated genes associated with bacterial growth, with down- and up-regulated genes indicated as bigger blue and red dots, respectively. (**C**) Score plot (PC2∼PC1) of the first (PC1) and second (PC2) principal components in principal component analysis (PCA) showed the clear separation between two transcriptome groups along the PC1 axis (accounting for 72.1% of total variance between specimens), revealing their different transcriptome profiles. (**D**) A volcano plot identified *E. faecalis* biofilm-associated differentially expressed genes (DEGs) (|log_2_(FC)| > 2.0 and P < 0.05), with down- and up-regulated genes labelled as blue and red dots, respectively. More stringent criteria (|log2(FC)| and -log(P) > 99% confidence intervals upper limits) were applied to identify the most significantly regulated genes associated with biofilm formation, with down- and up-regulated genes labelled as bigger blue and red dots, respectively. (**E** and **F**) Ten genes were identified as the most up-regulated genes that correlated with *E. faecalis* growth increase, based on more stringent criteria of both -log(P) (**E**) and log_2_(FC) (**F**). (**G** and **H**) Fifteen genes were identified as the most up-regulated genes that correlated with *E. faecalis* biofilm increase, based on more stringent criteria of both -log(P) (**G**) and log_2_(FC) (**H**). (**I** and **J**) Up-regulation of *E. faecalis fsrBD* at 9 hours (growth-promoting, **I**) and 48 hours (biofilm-promoting, **J**), that were induced by *K. pneumoniae*-produced active factors, were validated by RT-qPCR tests. (**K** and **L**) *E. faecalis* OG1RF grown in AUM supplemented with *K. pneumoniae* TOP52 supernatant (WT-Sup) gained enhanced growth (**K**) and biofilm formation (**L**), compared to grown in AUM only (WT-AUM), and deletion of genes *fsrBD* (Δ*fsrBD*-Sup) abolished this enhanced phenotype. Data represent the mean ± SD derived from three independent experiments. Statistics were performed using a two-sided unpaired t test with P ≤ 0.05 considered as statistically significant. *P ≤ 0.05, **P < 0.01, ***P < 0.001, ****P < 0.0001, ns indicates not significant.

To evaluate the effect of KpAUMSup on biofilm-related *E. faecalis* pathways, we took advantage of the finding that at 48 hrs post-inoculation, conditioned supernatant from *S. aureus* strain MRSA-1369 (SaAUMSup) stimulated *E. faecalis* growth in AUM to levels comparable to supplementation with KpAUMSup (∼2.81 logs increase) (fig. S10C), but unlike KpAUMSup, did not promote *E. faecalis* biofilm formation (fig. S10D). This suggested that a comparison between *E. faecalis* transcriptomes in AUM supplemented with KpAUMSup vs SaAUMSup would identify genes associated with biofilm. A PCA of an RNA-seq analysis conducted following 48 hrs of culture demonstrated distinct separation between KpAUMSup- vs SaAUMSup-derived transcriptomes along the PC1 axis (accounting for 72.1% of total variance) (Fig. 4C), indicating that these growth conditions produced distinct transcriptional profiles. The same stringent criteria (|log_2_(FC)| and - log(P) > 99% confidence intervals upper limits) identified the most significantly regulated biofilm-associated genes, including 27 up-regulated (Fig. 4, D, G, and H; table S11) and 33 down-regulated genes (Fig. 4D; table S11).

Among the two groups of up-regulated genes, only the *fsrBD* genes belonging to the *E. faecalis* Fsr quorum sensing system were identified with associations to both growth and biofilm (Fig. 5, E, F, G, and H). Quantitative reverse transcription polymerase chain reaction (RT-qPCR) tests validated up-regulation of *fsrBD* under conditions promoting growth (∼5.80 folds increase) (Fig. 4I) and biofilm (∼1.46 folds increase) (Fig. 4J). Further, an *fsrBD* deletion mutant (OG1RFΔ*fsrBD*) was attenuated for both planktonic growth (Fig. 4K) and biofilm (Fig. 4L) relative to wild-type strain OG1RF.

**Fig. 5.**
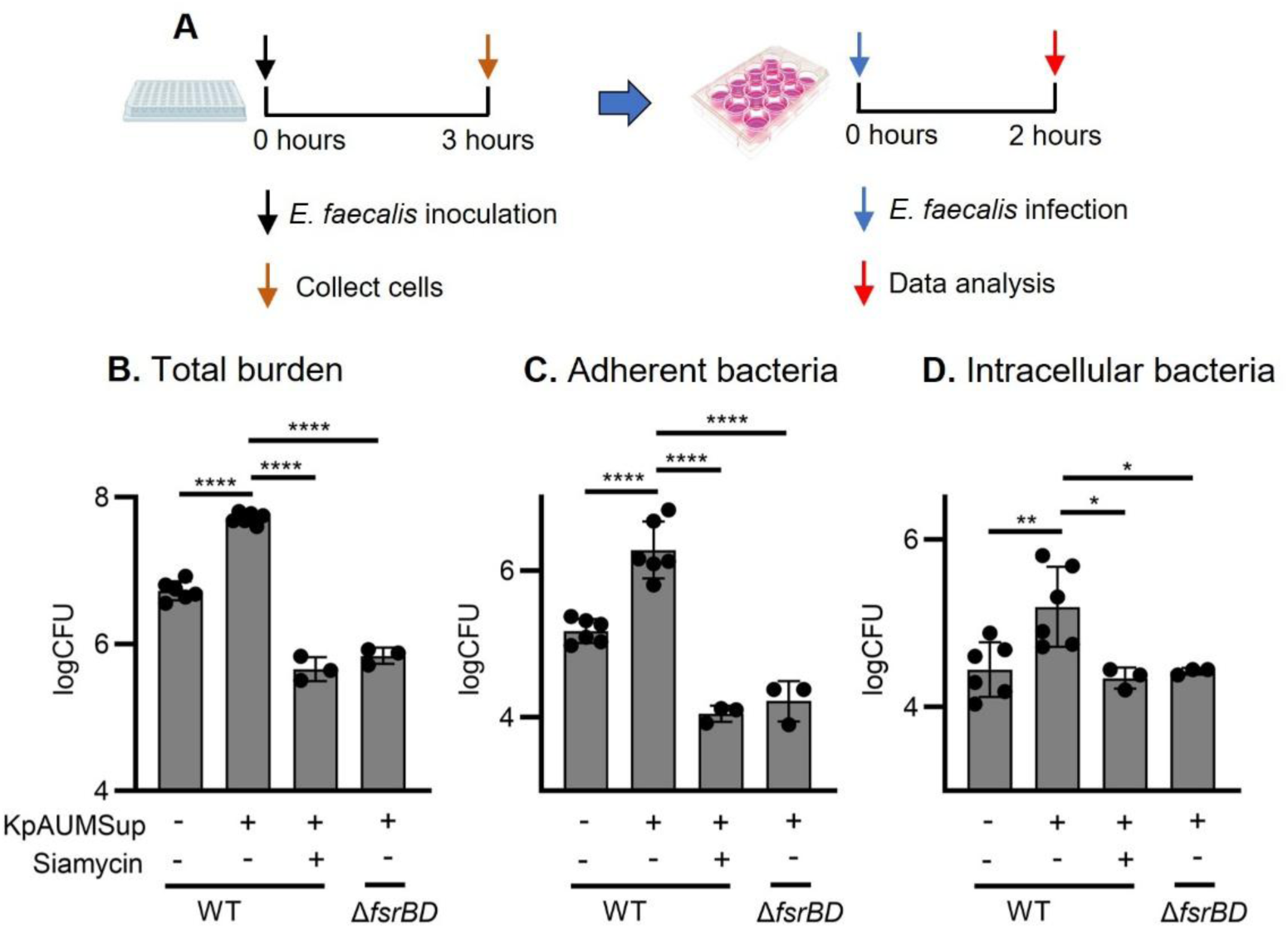
KpAUMSup enhances *E. faecalis* infection to human urothelial cells through a Fsr-dependent mechanism. Human urothelial 5637 cells were infected by *E. faecalis* OG1RF (WT) or an *fsrBD* deletion mutant (*ΔfsrBD*) following 3 hrs of pre-incubation in AUM, AUM supplemented with KpAUMSup with or without the Fsr inhibitor siamycin at a subinhibitory concentration (5.78 µM), as shown (**A**). Following 2 hrs of infection (**A**), cells were processed to determine (**B**) total bacterial burdens, (**C**) bacteria adherent to the host cell surface and (**D**) intracellular bacteria. Data represent the mean ± SD derived from at least three independent experiments. Statistics were performed using a two-sided unpaired t test with P ≤ 0.05 considered as statistically significant. *P ≤ 0.05, **P < 0.01, ***P < 0.001, ****P < 0.0001, ns indicates not significant.

### Fsr is required for planktonic growth and biofilm promoted by KpAUMSup

In the Fsr quorum sensing system, *fsrD* encodes the precursor of the autoregulatory peptide gelatinase biosynthesis-activating pheromone (GBAP) that is processed into its active form by FsrB (*18–20*). The extracellular concentration of GBAP is sensed by a two-component transcriptional regulator (encoded by *fsrA, fsrC*) to control multiple biofilm-, metabolism- and virulence-related genes, which most prominently include the secreted proteases GelE and SprE (*21–23*). While up-regulation was not observed in RNA-Seq datasets (table S12), a targeted examination by RT-qPCR revealed that both protease genes were up-regulated under conditions emphasizing growth (AUM vs KpAUMSup at 9 hrs post-inoculation) (fig. S11, A and C), but not conditions that emphasized biofilm (SaAUMSup vs KpAUMSup at 48 hrs post-inoculation) (fig. S11, B and D). Consistent with expression data, analysis of GelE and SprE deletion mutants (OG1RFΔ*gelE*, OG1RFΔ*sprE*) indicated that the loss of activity of either protease inhibited planktonic growth in the presence of KpAUMSup compared to WT OG1RF (OG1RFΔ*gelE* = ∼1.14 logs decrease, OG1RFΔ*sprE* = ∼1.12 logs decrease) (fig. S12A), but had no effect on biofilm (fig. S12B). In contrast, an *fsrBD* deletion mutant (OG1RFΔ*fsrBD*) was defective for both planktonic growth (∼2.13 logs decrease) and biofilm (∼2.24 folds decrease) under these conditions (fig. S12, A and B). Together, these data show that the ability of *E. faecalis* to respond to KpAUMSup with enhanced planktonic growth and biofilm requires Fsr quorum sensing, that Fsr-regulated protease activity contributes to growth, but not biofilm in urine-like conditions, suggesting that biofilm requires other Fsr-regulated pathways.

### KpAUMSup enhances *E. faecalis* infection of human urothelial cells through an Fsr-dependent mechanism

Infection of human urothelial 5637 cells *in vitro* was used to examine the effect of KpAUMSup on *E. faecalis* pathogenesis. Compared were *E. faecalis* OG1RF cultured under 3 conditions: i) AUM alone (OG1RF-AUM); ii) AUM supplemented with KpAUMSup (OG1RF-Sup); and iii) AUM + KpAUMSup with a subinhibitory concentration (5.78 µM) of the small molecule Fsr-inhibitor siamycin I (OG1RF-Siamycin) (*24*). An *fsrBD* deletion mutant (OG1RFΔ*fsrBD*) grown in AUM supplemented with KpAUMSup (Δ*fsrBD*-Sup) was also examined and compared. All cultures were harvested at 3 hrs post-inoculation for infection of 5637 cells (Fig. 5A, figs. S13 and S14). When analyzed following 2 hrs of infection (Fig. 5A), KpAUMSup significantly enhanced infection on the basis of measurement of total CFUs (∼0.98 logs increase) (Fig. 5B), numbers of bacteria adherent to cells (∼1.23 logs increase) (Fig. 5C) and intracellular CFUs (∼0.86 logs increase) (Fig. 5D) (Compare OG1RF-AUM to OG1RF-Sup). The loss of Fsr, through genetic deletion decreased total, adherent, and intracellular CFUs, by ∼1.87 logs, ∼2.16 logs, and ∼0.98 logs respectively. Chemical inhibition decreased total, adherent, and intracellular CFUs, by ∼2.04 logs, ∼2.37 logs, and ∼1.05 logs respectively. The loss of Fsr significantly reduced infection efficiency (compare OG1RF-AUM to Δ*fsrBD*-Sup, OG1RF-Siamycin, Fig. 5, B, C, and D). Together, these data demonstrate that KpAUMSup promotes enhanced infection of 5637 urothelial cells and the ability of *E. faecalis* to respond to KpAUMSup requires the Fsr quorum sensing system.

### KpAUMSup enhances infection in a murine model of CAUTI

To examine the effect of KpAUMSup on infection *in vivo*, we used the well-characterized murine CAUTI model (*25, 26*). Compared were *E. faecalis* WT OG1RF and the Fsr deletion mutant cultured under the conditions described above. Cultures were harvested following 9 hrs of incubation, a point just prior to when KpAUMSup initiates enhanced planktonic growth (see above, figs. S15 and S16). Groups of 7-week-old C57BL/6NCr mice received a transurethral catheter implant, were inoculated with 1 x 10^7^ CFU of bacteria and infection efficiency assessed following 5 hrs of infection (Fig. 6A). Similar to infection of 5637 urothelial cells, KpAUMSup significantly enhanced infection efficiency as determined by total CFUs recovered from the whole bladder (∼1.26 logs increase) (Fig. 6B) and the catheter implant (∼1.38 logs increase) (Compare OG1RF-AUM to OG1RF-Sup, Fig. 6, B and C). This enhanced efficiency for bladder and catheter CFUs was inhibited by both siamycin treatment (∼1.14 logs and ∼0.97 logs decreases, respectively) (OG1RF-Siamycin) and in the Fsr mutant (Δ*fsrBD*-Sup) (∼0.90 logs and ∼0.72 logs decreases for bladder and catheter CFUs, respectively) (Fig. 6, B and C). Together, these results demonstrate that KpAUMSup can enhance the ability of *E. faecalis* to infect the catheterized bladder through an Fsr-dependent mechanism.

**Fig. 6.**
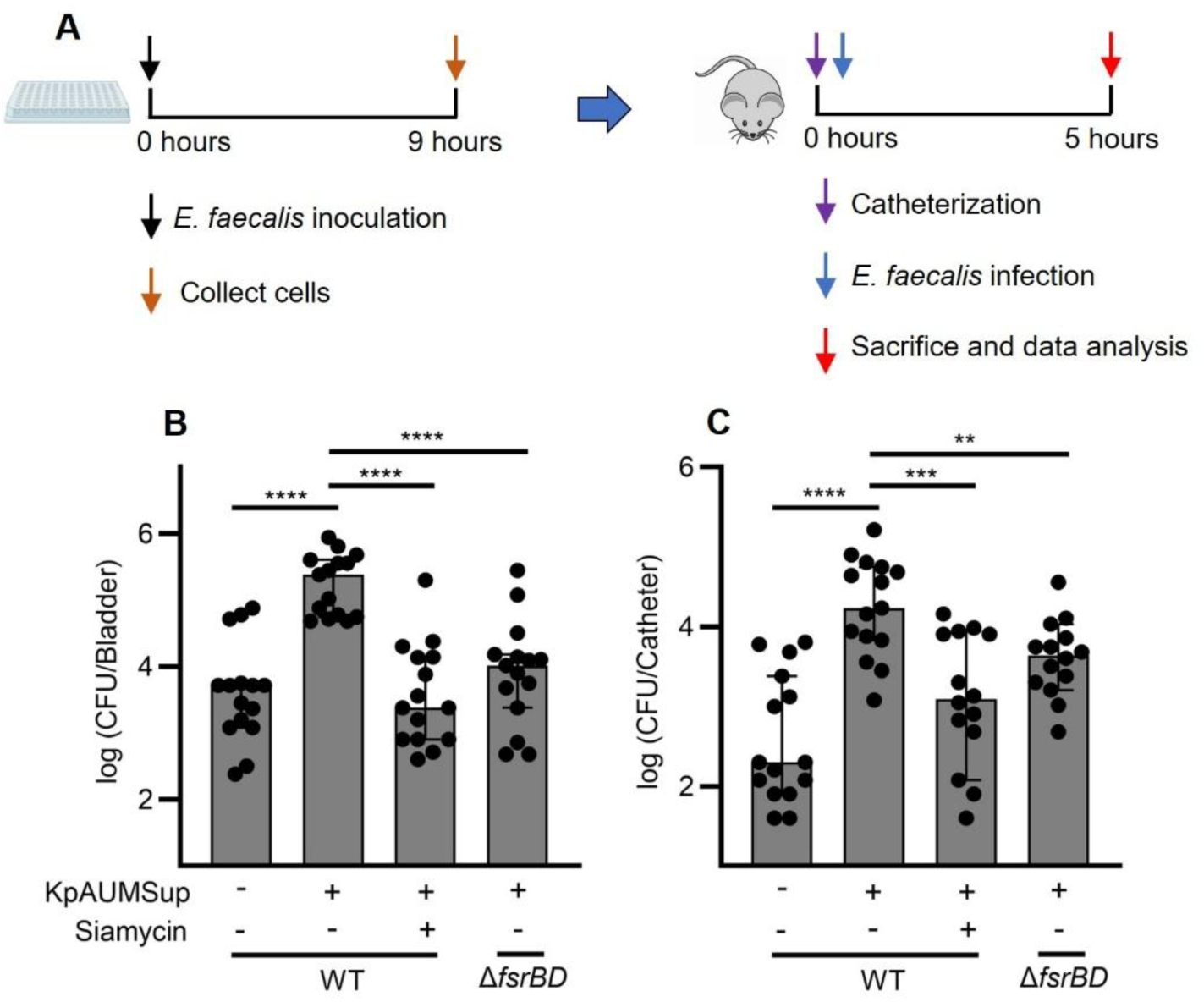
KpAUMSup enhances *E. faecalis* infection in a murine model of CAUTI. Seven-week-old female female C57BL/6NCr mice were transurethrally catheterized and infected by *E. faecalis* OG1RF (WT) or an *fsrBD* deletion mutant (*ΔfsrBD*) following 9 hrs of pre-incubation in AUM, AUM supplemented with KpAUMSup with or without the Fsr inhibitor siamycin at a subinhibitory concentration (5.78 µM), as shown (**A**). Following 5 hrs of infection (**A**), mice were euthanized and processed to determine the bacterial burdens in the (**B**) bladder and on the (**C**) catheter. Data represent the median ± 95% CI derived from at least two independent experiments. Statistics were performed Mann-Whitney U test with P ≤ 0.05 considered as statistically significant. *P ≤ 0.05, **P < 0.01, ***P < 0.001, ****P < 0.0001, ns indicates not significant.

## Discussion

Understanding the mechanisms that promote the formation of polymicrobial communities on urinary catheters leading to CAUTI will be essential for developing new therapeutic approaches to treat these typically multidrug-resistant infections. In the present study, we have shown that during culture in AUM, *K. pneumoniae* generates extracellular heat-sensitive small molecules that stimulate the Fsr quorum sensing system of *E. faecalis* to promote enterococcal planktonic growth and biofilm and to enhance efficiency of infection in several models of CAUTI. This beneficial interaction may help to explain both the high frequency of co-colonization of urinary catheters by these two pathogens and their resilience in long-term catheterized patients.

In the environment, multi-species microbial communities are typically challenged by the scarcity of essential nutrients, making trophic interactions a major driver of community assembly (*27, 28*). These interactions are defined by the exchange of metabolites between the members of a polymicrobial community and most commonly occur when a keystone species breaks down a complex nutrient into components that can be utilized by partner species or generates a by-product of its own metabolism that can promote the growth of other community members (*29–31*). Since urine is highly variable and nutrient limited (*32*), an important element of the current study is our use of AUM to more accurately reproduce conditions that exist in the urinary environment, which revealed an apparent trophic interaction of benefit to *E. faecalis* that was not evident in nutrient-rich medium. Our prior analyses have shown that *E. faecalis* can be restricted in its ability to grow in monoculture in human urine (*26*), indicating that our AUM culture system is of utility for analysis of complex trophic interactions in polymicrobial CAUTI. Our results suggest that *K. pneumoniae* with its larger genome and greater metabolic plasticity may be functioning as a keystone species to modify the bladder nutritional environment, either through the modification of a urine component or through a byproduct of its own metabolism. Which mechanism may be responsible for its beneficial interaction with *E. faecalis* will require the identification of the *K. pneumoniae* active factor.

Our data also indicate that the ability of the *K. pneumoniae* active factor to promote *E. faecalis* planktonic growth and biofilm is directly associated with its ability to stimulate the *E. faecalis* Fsr quorum sensing system. Considerable prior investigation has established that Fsr quorum-sensing plays a major role in regulating enterococcal metabolism, biofilm formation and virulence under nutrient-limited conditions (*18–20, 22, 23*). While it is possible that *K. pneumoniae* can produce a factor that directly stimulates the FsrC sensor kinase at low cell densities, inter-species quorum sensing typically involves the production of an autoinducer that is shared by both species, as exemplified by autoinducer-2 (AI-2), which is produced by numerous Gram-positive and - negative species (*33, 34*). In contrast, the autoinducer molecule of the Fsr system is a highly post-translationally modified peptide known as gelatinase biosynthesis-activating pheromone (GBAP) that is specific to *E. faecalis* (*35, 36*). Examination of the genomes of the *K. pneumoniae* strains used in this study did not reveal any polypeptide sequence with similarity to the GBAP pre-peptide, nor any enzyme similar to those required to generate the mature GBAP peptide, suggesting that stimulation of Fsr is through an indirect mechanism. Our data do show that an intact Fsr machinery is required, as deletion of the genes required for synthesis of GBAP (*fsrBD*) or siamycin inhibition of GBAP synthesis and sensing block the ability of the *K. pneumoniae* active factor to stimulate enterococcal planktonic growth and biofilm. Thus, while the mechanism of Fsr stimulation remains to be determined, these results implicate inter-bacterial signaling in promoting the positive association of *K. pneumoniae* and *E. faecalis* in polymicrobial CAUTI.

The canonical targets of Fsr quorum sensing are the secreted proteases Gelantinase (GelE) and Serine Protease (SprE), which have been shown to contribute to CAUTI in the murine model (*37, 38*). While neither of these proteases were found to be required for KpAUMSup to stimulate enterococcal biofilm in the *in vitro* AUM model used in the present study, the proteases are known to target the host in the CAUTI model, where SprE degrades regulators of the fibrinolytic pathway to increase the deposition of fibrinogen on the implanted catheter that serves a critical attachment protein that allows *E. faecalis* to form catheter biofilm in the bladder (*38*). Since our data show that Fsr is required for the ability of KpAUMSup to enhance virulence in the CAUTI model, stimulation of the Fsr-regulated proteases may contribute to this *in vivo* response. However, Fsr is known to regulate the expression of numerous other enterococcal genes involved in virulence, including several surface proteins involved in attaching to host cells or in biofilm formation under several different *in vitro* conditions (*39, 40*). Fsr also regulates genes involved in metabolism (*39*) which may contribute to the resiliency of *K. pneumoniae*-*E. faecalis* co-infection of catheters. Multiple studies have implicated bacterial doubling times in urine as a key variable that affects the outcome of bladder infection (*41, 42*), suggesting that during a catheter exchange in long-term catheterized patients, *K. pneumoniae* active factors provide *E. faecalis* with a growth advantage in the residual urine left after removal of an infected catheter, which allows it to outcompete other species for colonization of a newly inserted catheter.

Results of this study support a model of facultative mutualism where *K. pneumoniae* is acting as a keystone species to modify the bladder environment so that it can be exploited by *E. faecalis* through the production of a factor that actively simulates the Fsr regulatory pathway. However, the high rate of *K. pneumoniae*-*E. faecalis* co-occurrence on human catheters, their high frequency of re-establishing co-infection following catheter replacement and their high frequency of physical co-localization on human catheters (*16*) suggest a more interactive mutualistic relationship where *K. pneumoniae* also actively benefits from its interaction with *E. faecalis*. For example, there may be an additional trophic interaction where an *E. faecalis* metabolite promotes growth of *K. pneumoniae* in the bladder environment that was not captured in our AUM culture model. It has been shown that transport of ornithine from *E. faecalis* can stimulate arginine metabolism in *Proteus mirabilis* to facilitate polymicrobial biofilm and an increased incidence of the formation of urinary stones (*43*). Another possibility is that *K. pneumoniae* benefits from the ability of *E. faecalis* to suppress both innate and adaptive immunity (*44*). In this scenario, mutualism is promoted by the ability of *K. pneumoniae* to produce a factor stimulating Fsr to enhance *E. faecalis* growth, which in turn suppresses NF-kB signaling in innate immune cells to allow *K. pneumoniae* to evade host immunity (*45*).

The observation that many polymicrobial CAUTIs are composed of pairs of diverse bacterial species that are associated with each other at rates greater or less than that expected by chance alone, suggests that there are active mechanisms that support the assembly of these communities. The current study reveals a metabolic cooperation between *E. faecalis* and *K. pneumoniae* that facilitates their resilient co-infection, and demonstrates the considerable promise of developing targeted strategies for inhibiting key bacterial pathways to prevent positive interspecies interactions. Future work can be expanded to investigate diverse CAUTI pathogens for identifying pathogen-specific interaction-associated pathways to better elucidate the pathogenesis mechanism of polymicrobial CAUTI. Our findings lay the groundwork for developing novel therapeutics to perturb the polymicrobial community to prevent CAUTI.

## Methods

### Bacterial strains

*E. faecalis* and *K. pneumoniae* clinical isolates were collected from two long-term catheterized patients, Patients 85 and 82, from our previous clinical study (*16*). Prototypical reference strains *E. faecalis* OG1RF, *K. pneumoniae* TOP52, and MRSA-1369 were also used as described (*26, 46, 47*). *E. faecalis* OG1RFΔ*fsrBD*, OG1RFΔ*gelE*, and OG1RFΔ*sprE* are in-frame deletion mutants generated in a prior study (table S13) (*37*).

### Whole-genome sequencing

Bacterial genomic DNAs from 40 clinical and two prototypical reference strains, including 11 *E. faecalis* and 11 *K. pneumoniae* isolates from Patient 85, 9 *E. faecalis* and 9 *K. pneumoniae* isolates from Patient 82, and *E. faecalis* OG1RF and *K. pneumoniae* TOP52 strains, were extracted using a DNeasy PowerBiofilm Kit (Qiagen, 24000-50) from 1 mL overnight cultures. In total, 5 ng of DNA was used as input for library preparation and sequencing using a NovaSeq 6000 system (Illumina), following the protocol as previously described (*17*). The reads were demultiplexed by barcode, had adapter sequences removed with trimmomatic v.38, assembled into draft genomes using SPAdes v3.12.0 (Bankevich), and annotated for protein coding sequences on all contigs > 500 bp using prokka v1.12 (*17*). Phylogenetic analysis was performed by constructing core-genome alignment into a maximum likelihood tree using raxML and viewed in iTOL (*48*). Antibiotic resistant genes (ARGs) were identified by protein BLAST comparison to the CARD based on stringent cutoffs (> 95% identity and > 95% overlap with subject sequence) (*17, 49*). Average nucleotide identity (ANI) value was calculated using the EzBioCloud platform (*50*) by comparing genomic similarity between two strains among the four groups of clinical isolates from Patients 85 and 82, with the first isolate serving as the reference genome for comparison to all subsequent isolates. Isolates were considered to be the same species or the same or a nearly identical strain for ANI ≥ 95% and ANI ≥ 99.5%, respectively (*51, 52*).

### Culture and quantification

Bacterial growth and biofilm formation of paired *E. faecalis* and *K. pneumoniae* clinical isolates from Patients 85 and 82, and reference strains *E. faecalis* OG1RF and *K. pneumoniae* TOP52, were examined in microplate cultures. In brief, *E. faecalis* and *K. pneumoniae* overnight cell cultures were harvested by centrifugation, washed three times with an equal volume of PBS, and resuspended in an equal volume of fresh PBS solution. The bacterial suspension was used to inoculate fresh AUM (*17*) or BHI (Becton Dickinson, 237200) media at a dilution of 1:1000, with one species inoculated for monoculture or two species inoculated at 1:1 ratio for co-culture, followed by dispensing 200 µL aliquots into the wells of a 96-well plate. At 48 hours post-inoculation, bacterial growth and biofilm formation were quantified by CFU enumeration and crystal violet (CV) staining, respectively, using the methods as previously described (*17, 53*).

For testing biofilm formation on a catheter surface, a 1 cm platinum-cured silicone urinary catheter (Nalgene 50, 8060-0020) was cut into two equal pieces, one piece was added into a well containing 2mL of indicated bacterial culture in a 12-well plate. The plate was incubated at 37°C for 48 hrs followed by planktonic and sessile bacteria quantified by determination of CFUs as described (*17*). In all assays, *E. faecalis* strains were resistant to Kanamycin (50 μg/mL) but sensitive to Vancomycin (6 μg/mL), while *K. pneumoniae* strains were resistant to Vancomycin (6 μg/mL) but sensitive to Kanamycin (50 μg/mL), therefore plating on LB agar plates containing these antibiotics were used for differential enumeration of species-specific CFUs, as described^16^.

### Culture with conditioned medium supernatant

*E. faecalis* growth and biofilm formation in AUM were also examined in cultures supplemented with *K. pneumoniae* or MRSA-1369 conditioned AUM supernatant. *K. pneumoniae* or MRSA1369 strains were first cultured in AUM in individual wells of a 96-well microplate for 24 hours, growth from wells was pooled, and then filtered (Millipore Sigma, SLGPR33RB) to remove bacterial cells (*17, 54, 55*). The resulting supernatants (SaAUMSup, KpAUMSup) were then lyophilized and used to supplement o fresh *E. faecalis* AUM at a ratio of 1:1 for culture in microplate and catheter assays.

### Scanning electron microscopy

For each experiment, a single 1 cm platinum-cured silicone urinary catheter (Nalgene 50, 8060-0020) was cut into four equal pieces, and each piece was placed into an individual well of a 12-well plate containing 2 mL culture medium under three different conditions, including: i.) *E. faecalis* OG1RF in AUM, ii.) *E. faecalis* OG1RF and *K. pneumoniae* TOP52 in AUM, and iii.) *E. faecalis* OG1RF in AUM with KpAUMSup. At 48 hrs post-inoculation, each catheter segment was-t removed, was washed once with an equal volume of PBS, and then fixed with a solution of 2.5% glutaraldehyde/2% paraformaldehyde in 0.15M cacodylate buffer (pH = 7.4) for 15 min at room temperature, followed by further fixation in the same solution for overnight at 4°C. Samples were adhered to a conductive carbon adhesive tab on an aluminum stub, sputter coated with 6-nm iridium (Leica ACE 600) and imaged on a high-resolution FE-SEM system (Zeiss Merlin equipped with a Gemini II electron column) as previously described (*47*).

### Human catheter immunofluorescence imaging

Human catheters from Patients 85 and 82 were fixed in formalin and prepared for staining and imaging as previously described (*16, 56, 57*). In brief, catheter samples were washed three times with PBS, blocked with PBS containing 1.5% BSA and 0.1% sodium azide overnight at 4°C, and washed three times with PBS-T (1×PBS, 0.1% w/v Tween 20). Catheters were then stained with anti-*Enterococcus* polyclonal antibody (PA1-73120, Invitrogen) and anti-*Klebsiella* monoclonal antibody (Invitrogen, MA1-83282) at 1:400 dilution in dilution buffer (PBS-T, 0.1% w/v BSA, 0.5%methyl alpha-Dmannopyranoside) for 2 hours at room temperature, then washed and stained with IRDye 680LT Donkey anti- Rabbit (LI-COR, 926-68023) and IRDye 800CW Donkey anti-Mouse (LICOR, 926-32212) antibodies at 1:1000 dilution for 1 hour at room temperature. To determine background levels of fluorescence, control catheters were stained with the secondary fluorescent antibodies in the absence of the primary antibodies. Finally, catheters were washed with PBS-T three times and visualized using an Odyssey Imaging System (LI-COR Biosciences).

### Biofilm immunofluorescence imaging

*E. faecalis* OG1RF cultures, in AUM or AUM supplemented with KpAUMSup from TOP52, were prepared using 5 mL of medium in a 35 mm diameter culture dish (MatTek, P35G-0-14-C). At 48 hrs post-inoculation, media were removed, each well was washed with PBS, fixed by the addition of 10% neutral buffered formalin solution (Millipore Sigma, HT501128). Following incubation overnight at 4°C, wells were stained with anti-*Enterococcus* polyclonal primary antibody (Invitrogen, PA1-73120) at 1:500 dilution overnight at 4°C, followed by anti-rabbit IgG cross-adsorbed secondary antibody (Invitrogen, A-11008) at 1:500 dilution for 3 hrs at room temperature, and imaged using a Zeiss LSM 880 Confocal Laser Scanning Microscope, with biofilm volume quantified using Volocity software (Quorum) as described (*58, 59*).

### RNA Sequencing and comparative transcriptomic analysis

RNA sequencing (RNA-seq) and comparative transcriptomic analysis were used to compare: i) *E. faecalis* OG1RF grown in AUM in the presence or absence of KpAUMSup from TOP52 conditioned supernatant for 9 hrs to identify growth-associated differentially regulated genes, and ii) *E. faecalis* OG1RF grown in AUM supplemented with SaAUMSup from MRSA-1369 or TOP52 KpAUMSup for 48 hrs to identify biofilm-associated differentially regulated genes. Briefly, *E. faecalis* OG1RF microplate cultures were grown as described above, multiple wells (200 µL/well × 50 wells = 10 mL total culture) were harvested, pooled, and subjected to centrifugation to collect bacterial pellets. RNA was extracted from bacterial pellets using the RNeasy Plus Mini Kit (Qiagen, 74134) with the quality of the purified RNA determined by spectroscopy (NanoDrop 2000, Thermo Fisher). Libraries for Illumina sequencing were prepared using the FastSelect RNA kit (Qiagen, 334222), according to the manufacturer’s protocol and sequences were determined using an Illumina NovaSeq 6000 and processed as previously described to generate RNA-seq reads for comparative transcriptomic analysis (*53, 60–67*). Comparative transcriptomic analyses, including principal component analysis (PCA) for identifying transcriptomes differences and differential expression analysis for determining differentially expressed genes, were conducted as previously described (*17, 53*).

### Quantitative reverse transcription polymerase chain reaction (RT-qPCR)

Transcription levels of *E. faecalis* OG1RF genes *gyrA*, *fsrBD*, *gelE*, and *sprE* under both growth-and biofilm-promoting conditions were examined by quantitative reverse transcription polymerase chain reaction (RT-qPCR) test. RNA was extracted and prepared as described above. Reverse transcription, polymerase chain reaction (PCR) amplification, and real-time PCR (qPCR) were performed using specific primers (Table 1) following the protocols as previously described (*53*). Each sample was run in triplicate with average Ct values calculated, followed by relative expression compared to control determined by the ΔΔCt method (*53, 68*).

**Table 1.**
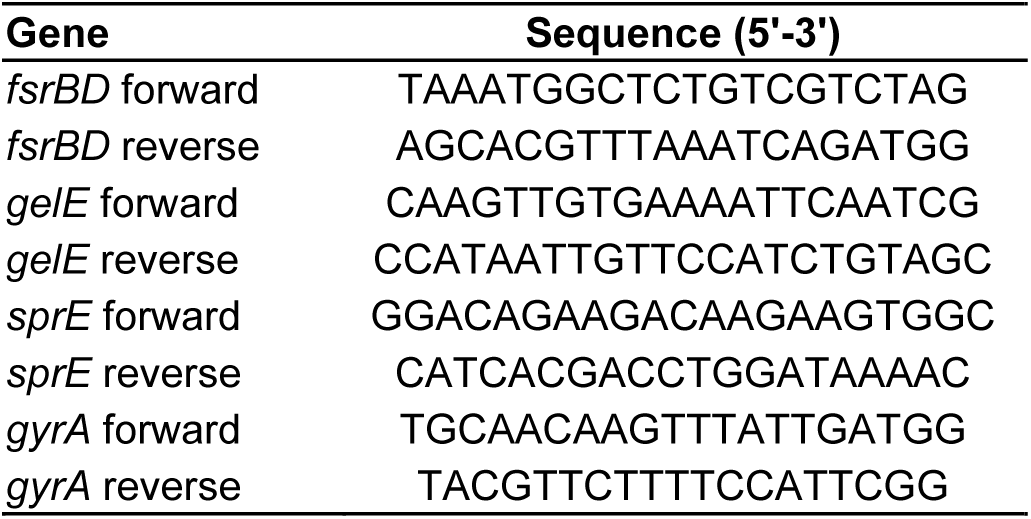
Primers list.

### Fractionation of KpAUMSup by filtration

Separation of KpAUMSup into fractions into low and high molecular weight (MW) fractions was conducted as follows: *K. pneumoniae* TOP52 was cultured overnight in AUM and bacterial cells removed by centrifugation and filtration as described above. The resulting supernatant was added to an ultracentrifuge filter tube with a 5000 molecular weight cut-off (MWCO) (Tisch Scientific), and subject to centrifugation at 7000g for 2 hrs (Eppendorf 5430R). Both the hold up (<5000 Da) and flow-through (>5000 Da) fractions were collected, lyophilized, and used to supplement *E. faecalis* AUM cultures as described above.

### High-performance liquid chromatography (HPLC) fractionation of *K. pneumoniae* supernatant

*K. pneumoniae* TOP52 AUM supernatant was collected using the method as described above, followed by fractionation via two consecutive steps, including an Amberlite XAD16N resin (20∼60 mesh; Sigma) and an Ascentis Express Phenyl-Hexyl column (150 × 4.6 mm, 2.7 μm; Sigma–Aldrich) for purifying and concentrating *K. pneumoniae*-produced active factors. In brief, culture supernatant was first incubated with a methanol (100%)-conditioned Amberlite resin (Sigma) for 24 hrs and eluted with 100% methanol. The eluate was concentrated in a rotatory evaporator (R-100 Rotavapor; BUCHI), lyophilized (Labconco), resuspended in HPLC-grade water plus 0.1% formic acid, and further purified on a Bio-Rad BioLogic DuoFlow 10 system equipped with a QuadTec UV–Vis detector and an Ascentis Express Phenyl-Hexyl column (Sigma–Aldrich) (*69*). The Ascentis Express Phenyl-Hexyl column was run at 0.30 ml/ min with HPLC-grade water plus 0.1% formic acid (solvent A) and acetonitrile plus 0.1% formic acid (solvent B) using a gradient as follows. Solvent B held steady at 2% for 1.0 ml, then increased to 95% over 40 ml, and finally increased to 100% over 1 ml. Elute collection started following 1mL and a series of 2 ml fractions were collected for a total of 20 fractions. Individual HPLC fractions were lyophilized and used to supplement *E. faecalis* AUM cultures at a volume ratio of 1:1 in the microplate format to determine fractions with the ability to enhance *E. faecalis* growth and biofilm formation, as described above.

### *E. faecalis* culture with heat-treated *K. pneumoniae* factors

To examine the sensitivity of *K. pneumoniae*-produced factors to heat treatment, both *K. pneumoniae* TOP52 AUM supernatant and functional fractions obtained from supernatant fractionation as described above, were contained in sealed conical tubes (Labcon) and heated at 95°C in water bath (Thermo Scientific Precision 280) for 24 hrs, filtered, lyophilized, and added to fresh *E. faecalis* OG1RF AUM cultures to test their activities to promote growth and biofilm formation of *E. faecalis* using the methods described above.

### Determination of subinhibitory siamycin concentrations

The Fsr inhibitor siamycin (*24*) was used at the highest concentration that did not inhibit planktonic growth determined as follows. Various concentrations of siamycin I (Cayman Chemical) were added fresh AUM supplemented with TOP52 KpAUMSup, which were then inoculated with *E. faecalis* OG1RF and cultured at 37°C for 3 or 9 hrs for determination of CFUs. The siamycin concentration used for subsequent experiments was the highest concentration that inhibited enterococcal CFUs recovered by <5% relative to untreated cultures.

### Infection of human urothelial cells

Human urothelial 5637 cells were cultured as previously described (*70, 71*). In brief, 2 mL of 10^5^ cells/mL 5637 urothelial cells in RPMI medium (Gibco) plus 10% heat inactivated fetal bovine serum (HI-FBS, Omega) were seeded to each well of a 12-well tissue culture plate (TPP Techno, Z707775) and incubated at 37°C in an atmosphere of 5% CO_2_ (Forma Series II Incubator, Thermo) overnight to reach approximately 80% confluency. Cultures in microplates were prepared and started as described above, including i.) *E. faecalis* OG1RF grown in AUM, ii.) *E. faecalis* OG1RF grown in AUM supplemented with TOP52 KpAUMSup, iii.) *E. faecalis* grown in AUM supplemented with TOP52 KpAUMSup plus a subinhibitory concentration of siamycin I (5.78 µM), and iv.) *E. faecalis* OG1RFΔ*fsrBD* grown in AUM supplemented with TOP52 KpAUMSup. At 3 hrs post-inoculation, the cells were harvested by centrifugation, resuspended in RPMI medium plus 10% HI-FBS, and added to 5637 monolayers (1 mL/well, ∼10^5.5^ CFU/mL). Following 2 hrs of infection at 37°C, CFUs were determined, including: i.) total CFUs (extracellular, adherent and intracellular bacteria) ii.) bacteria adherent to cells and iii.) intracellular bacteria. To quantify total CFUs, after 2 hrs of incubation, 20 μL 5% Triton X-100 (Millipore Sigma, X100) was added and incubated at 37°C for 45 min to lyse cells, followed by scraping wells and CFU enumeration by quantitative plating. To quantify intracellular bacteria, after 2 hrs of incubation, the cells were washed twice with an equal volume of PBS, treated with Vancomycin (100 μg/mL) at 37°C for 2 hrs, washed again for three times with an equal volume of PBS, lysed with 1 mL of 0.1% Triton X-100 at 37°C for 45 min, followed by scraping and CFU enumeration. To quantify bacteria adherent to the host cell surface, after 2 hrs of incubation, the cells were washed for five times with an equal volume of PBS, lysed with 1 mL of 0.1% Triton X-100 at 37°C for 45 min, followed by scraping and CFU enumeration which quantified the sum of adherent and intracellular bacteria. Finally, the amount of bacteria adherent to the host cell surface was obtained by subtracting the amount of intracellular bacteria from the sum of adherent and intracellular bacteria.

### Mouse catheter implantation and infection

For preparation of the inoculum, WT *E. faecalis* OG1RF and OG1RFΔ*fsrBD* were cultured in microplates under the several conditions described above, harvested at 9 hrs post-inoculation and prepared as described above. Catheter implantation and *E. faecalis* infection were conducted as previously described (*25*). In brief, seven-week-old C57BL/6NCr mice (Charles River Laboratories) were transurethrally catheterized with a segment of 4∼5 mm silicone tubing and immediately infected transurethrally with a 50 µL volume containing ∼1 × 10^7^ CFU of bacteria suspended in PBS Mice were euthanized at 5 hrs post-infection, bladders and catheters were collected and processed for determining bacterial CFUs as previously described (*25*). All animal experimentation was conducted following the National Institutes of Health guidelines for housing and care of laboratory animals and performed in accordance with institutional regulations under the supervision of the Animal Studies Committee at Washington University School of Medicine (Protocol 24-0279, Animal Welfare Assurance #D16-00245).

### Statistics

Unless otherwise stated, conclusions were based on the comparison of means or medians generated from at least three biological replicates, each of which were tested in at least two independent experiments, that were tested for significance using a two-tailed unpaired *t* test or a Mann Whitney *U* test and conducted using GraphPad Prism 9.4.1 (GraphPad software). P ≤ 0.05 was considered significant.

### Data and code availability

The genomes and RNA-Seq reads analyzed in this study have been deposited to NCBI database under the project accession no. PRJNA1136145 and no. PRJNA1140253, respectively. The computer codes for the analyses in this study are available in Github (https://github.com/QL5001/PolyEK_script; branch name, main; commit ID, a62d866).

## Acknowledgements

We thank Wandy Beatty for assistance with confocal microscopy. We thank Washington University Center for Cellular Imaging (WUCCI) for assistance with scanning electron microscopy. This work was supported by the National Institutes of Health grant RO1DK51406 (SJH and MGC). It was also supported by the Centers for Disease Control Prevention Epicenters Program Grant (CU54 CK 000162) (JPH) and the National Institutes of Health grants R01DK099534 (JPH) and R01DK111930 (JPH).

## Author contributions

Conceptualization: ZZ, JSP, KWD, JPH, SJH, MGC. Methodology: ZZ, JSP, KWD, JPH, SJH, MGC. Investigation: ZZ, JSP, CLPO, KRK, DAS, SMH. Visualization: ZZ, JPH, SJH, MGC. Validation: ZZ, JSP, JPH, SJH, MGC. Writing—original draft: ZZ, SJH, MGC. Writing—review & editing: ZZ, KWD, JPH, SJH, MGC.

