## Supplementary Materials for "Metabolic cross-talk promotes persistence of *Enterococcus* in a model of polymicrobial catheter-associated urinary tract infection"

**Table of contents for supplementary materials**

Figs. S1 to S16

Tables S1 to S13

### SUPPLEMENTARY FIGURES

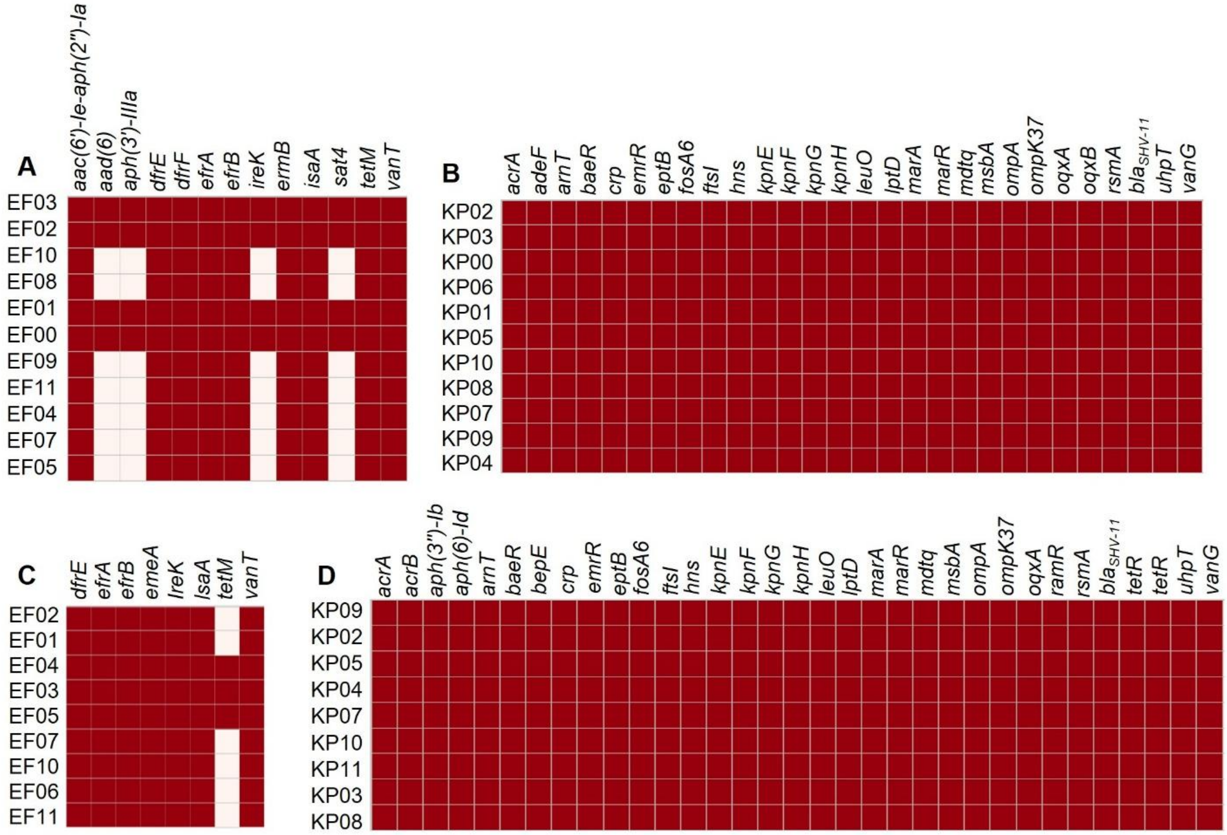


**Fig. S1.** ***E. faecalis*** **and** ***K. pneumoniae*** **strains cause recurrent CAUTI in long-term catheterized patients.** Antibiotic resistance genes (ARGs) identifications of 11 *E. faecalis* (**A**) and 11 *K. pneumoniae* (**B**) clinical isolates from Patient 85, and 9 *E. faecalis* (**C**) and 9 *K. pneumoniae* (**D**) clinical isolates from Patient 82.

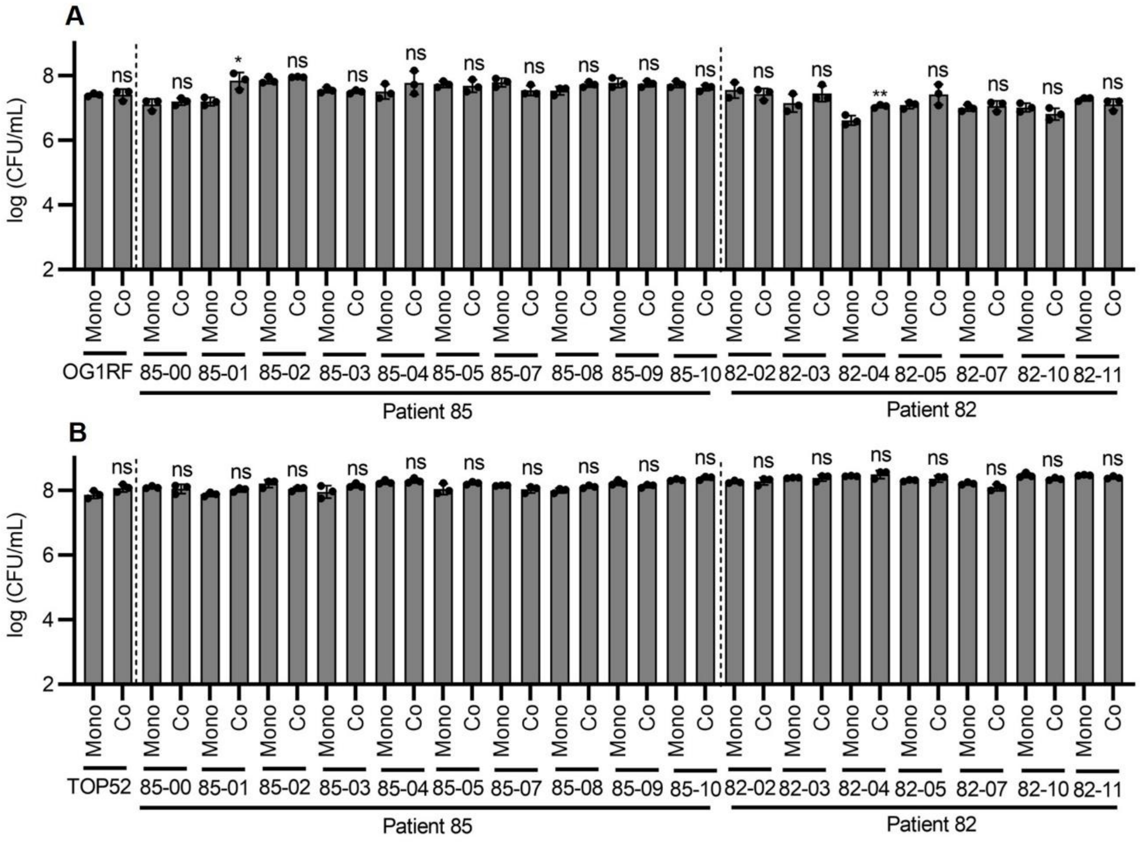


**Fig. S2. *K. pneumoniae* did not promote *E. faecalis* growth in co-cultures using nutrient-sufficient BHI media.** (**A** and **B**) Growth of clinical *E. faecalis* (**A**) and *K. pneumoniae* (**B**) isolates collected at the same collection periods from Patients 85 and 82 in monocultures (Mono) and mixed cultures (combo) in AUM in microplate assays. Prototypical *E. faecalis* OG1RF and *K. pneumoniae* TOP52 strains were included. Data represent the mean ± SD derived from three independent experiments. Statistics were performed using a two-sided unpaired t test with $P \leq 0.05$ considered as statistically significant. *$P \leq 0.05$, **$P < 0.01$, ***$P < 0.001$, ****$P < 0.0001$, ns indicates not significant.

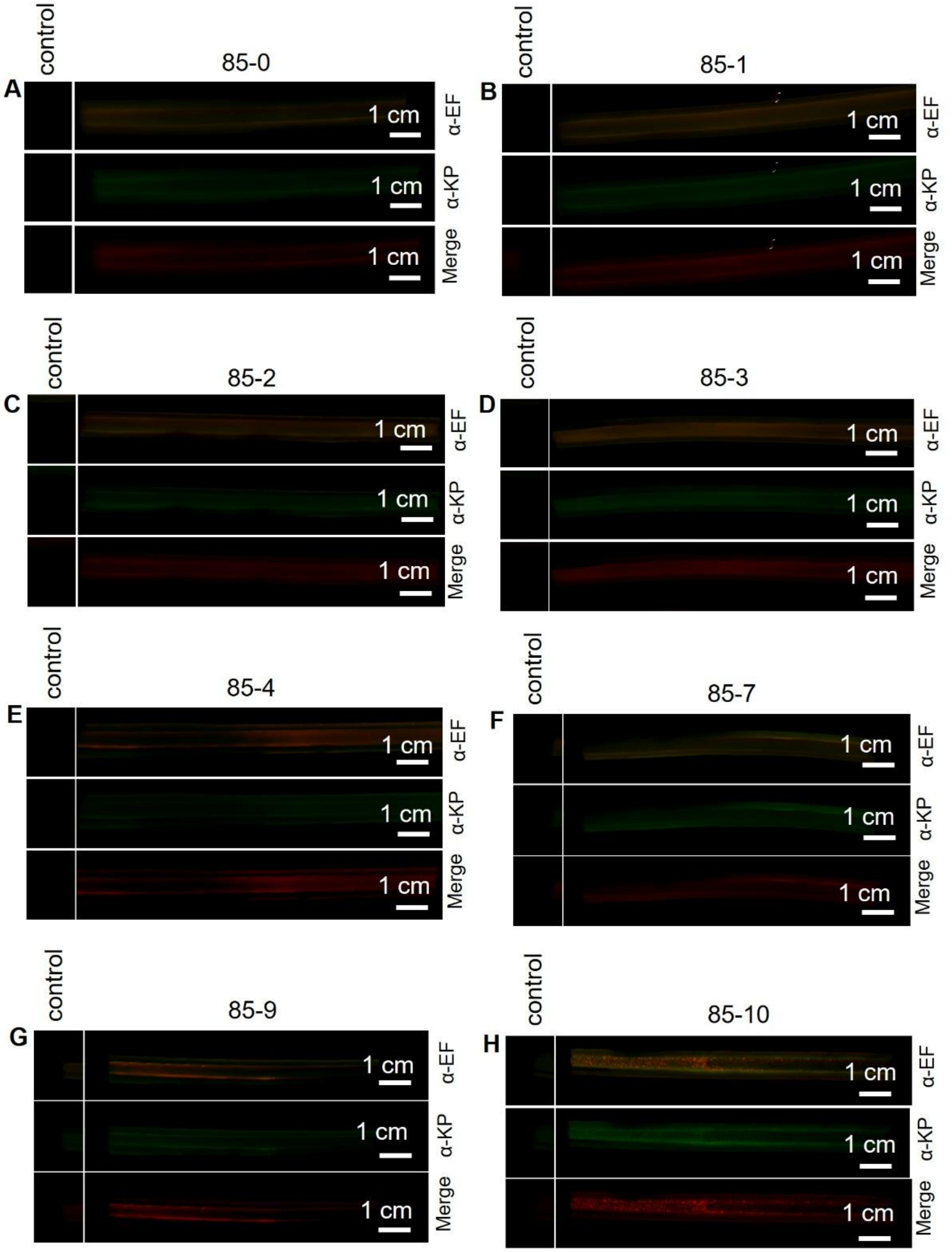


**Fig. S3.** Immunohistochemistry imaging of clinical *E. faecalis* and *K. pneumoniae* isolates on formalin-fixed catheter portions collected at 8 indicated periods from Patient 85, including period 0 (**A**), period 1 (**B**), period 2 (**C**), period 3 (**D**), period 4 (**E**), period 7 (**F**), period 9 (**G**), period 10 (**H**).

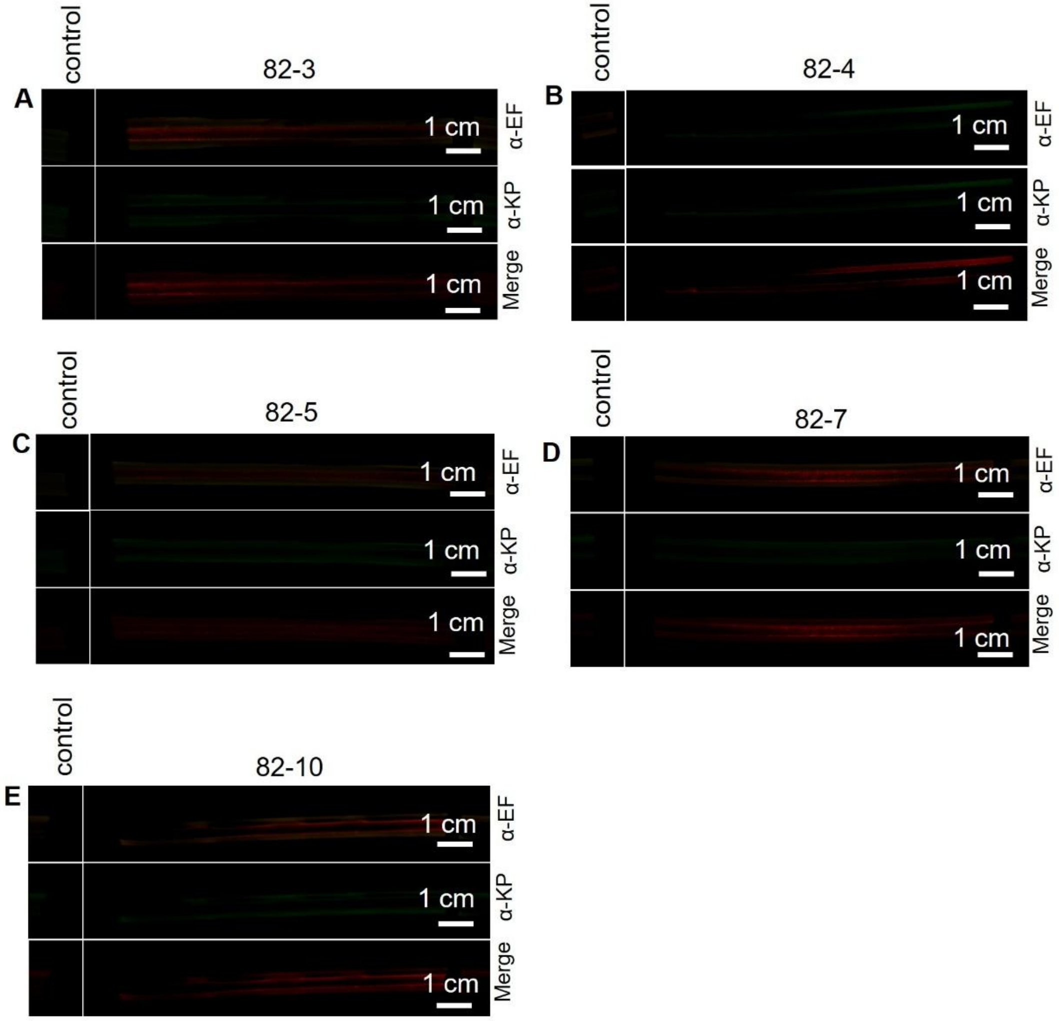


**Fig. S4.** Immunohistochemistry imaging of clinical *E. faecalis* and *K. pneumoniae* isolates on formalin-fixed catheter portions collected at 5 indicated periods from Patients 82, including period 3 (**A**), period 4 (**B**), period 5 (**C**), period 7 (**D**), period 10 (**E**).

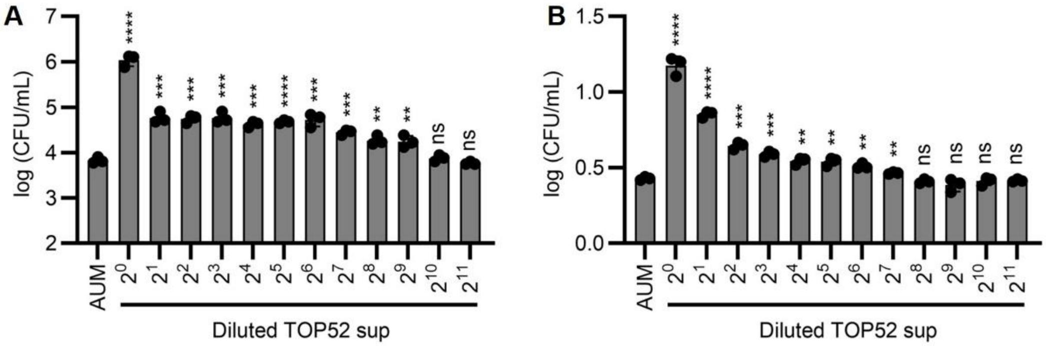


**Fig. S5. *K. pneumoniae*-generated active factors promote *E. faecalis* growth and biofilm formation.** (**A** and **B**) Bacterial growth (**A**) and biofilm formation (**B**) of *E. faecalis* OG1RF grown in AUM (AUM) and AUM supplemented with serially diluted *K. pneumoniae* TOP52 supernatant (from $2^0$ to $2^{11}$ dilutions) in microplate assays. Data represent the mean ± SD derived from three independent experiments. Statistics were performed using a two-tailed unpaired t test with $P \leq 0.05$ considered as statistically significant. *$P \leq 0.05$, **$P < 0.01$, ***$P < 0.001$, ****$P < 0.0001$, ns indicates not significant.

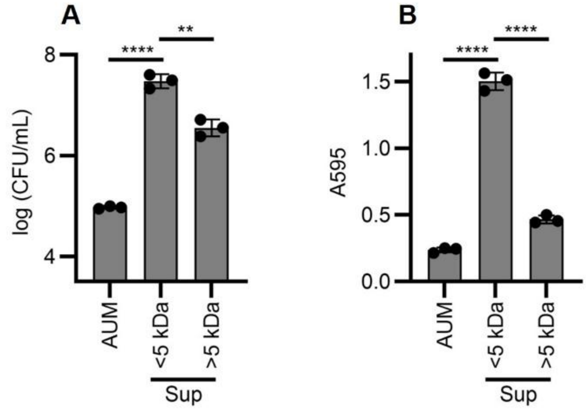


**Fig. S6. *K. pneumoniae*-produced active factors are smaller than 5 kDa.** (**A** and **B**) Bacterial growth (**A**) and biofilm formation (**B**) of *E. faecalis* OG1RF in only AUM (AUM), and in AUM supplemented with *K. pneumoniae* TOP52-produced extracellular compounds whose molecular weight were smaller (< 5 kDa) and bigger (> 5 kDa) than 5 kDa. Data represent the mean ± SD derived from three independent experiments. Statistics were performed using a two-tailed unpaired t test with $P \leq 0.05$ considered as statistically significant. *$P \leq 0.05$, **$P < 0.01$, ***$P < 0.001$, ****$P < 0.0001$, ns indicates not significant.

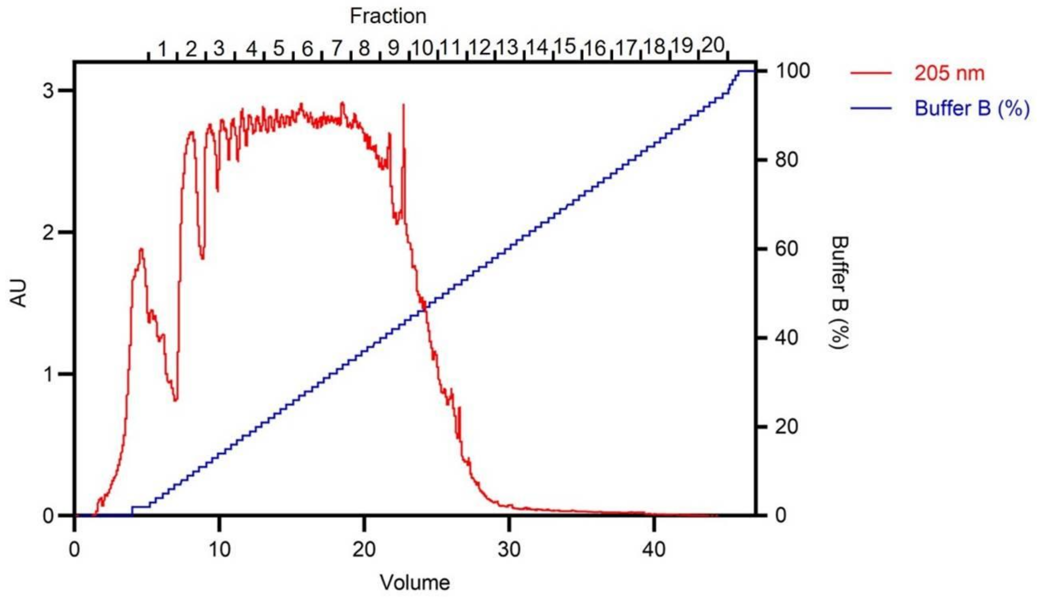


**Fig. S7. High-performance liquid chromatography fractionation of *K. pneumoniae* supernatant.** *K. pneumoniae* TOP52-conditioned AUM culture supernatant was first fractionated by Amberlite resin, with the elute purified by a reverse-phase HPLC fractionation which yielded 20 fractions.

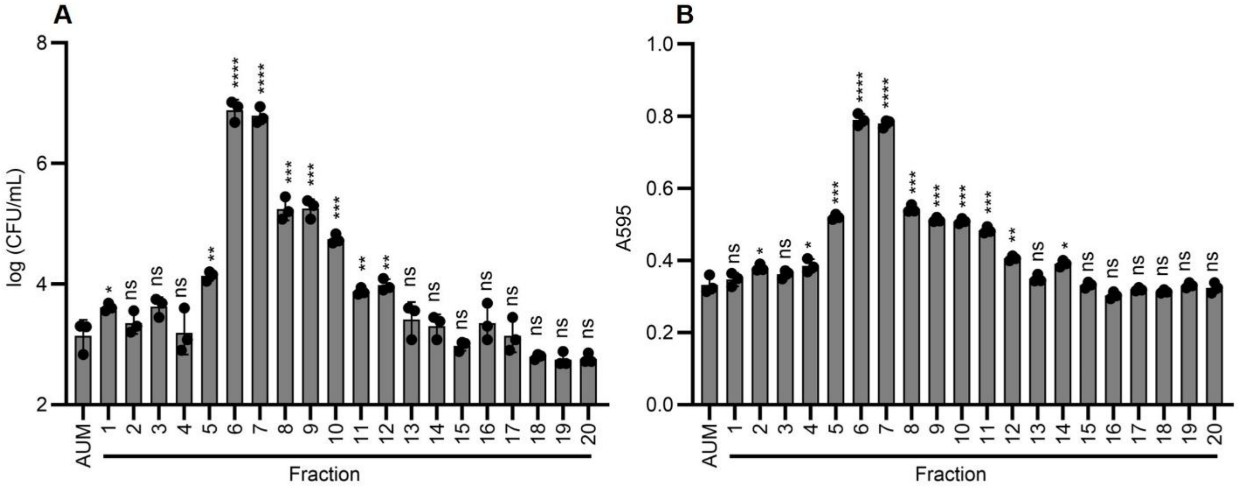


**Fig. S8. Functional fractions promote enhanced *E. faecalis* growth and biofilm formation.** (**A** and **B**) Two fractions, Fraction 6 and 7, collected from HPLC fractionation of *K. pneumoniae* TOP52 AUM supernatant were identified with the activities to enhance bacterial growth (**A**) and biofilm formation (**B**) of *E. faecalis* OG1RF. Data represent the mean ± SD derived from three independent experiments. Statistics were performed using a two-sided unpaired t test with $P \leq 0.05$ considered as statistically significant. $^{*}P \leq 0.05$, $^{**}P < 0.01$, $^{***}P < 0.001$, $^{****}P < 0.0001$, ns indicates not significant.

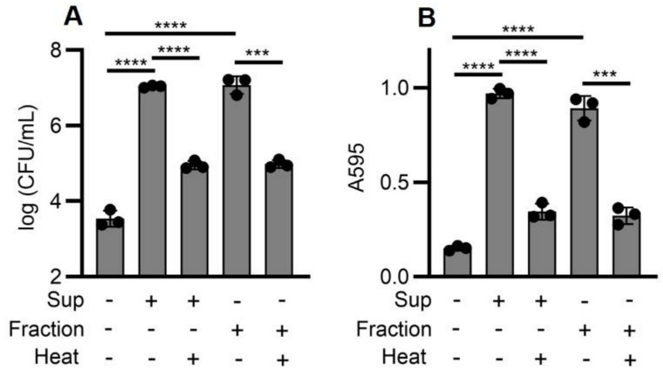


**Fig. S9. *K. pneumoniae*-produced active factors are identified with heat-labile properties.** (**A** and **B**) Bacterial growth (**A**) and biofilm formation (**B**) of *E. faecalis* OG1RF in AUM only, in AUM supplemented with *K. pneumoniae* TOP52 supernatant without or with heating treatment, and in AUM supplemented with active HPLC fractions without or with heating treatment. Data represent the mean ± SD derived from three independent experiments. Statistics were performed using a two-sided unpaired t test with $P \leq 0.05$ considered as statistically significant. *$P \leq 0.05$, **$P < 0.01$, ***$P < 0.001$, ****$P < 0.0001$, ns indicates not significant.

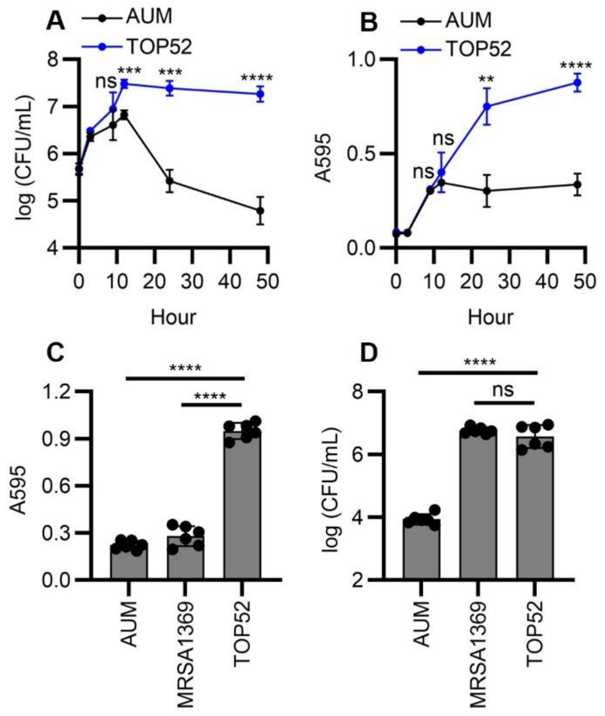


**Fig. S10. *E. faecalis* growth and biofilm formation are affected by extracellular factors produced by *K. pneumoniae* .** (**A** and **B**) Curves of bacterial growth (**A**) and biofilm formation (**B**) for *E. faecalis* OG1RF grown in AUM (AUM) and AUM supplemented with *K. pneumoniae* TOP52 AUM culture supernatant (+TOP52Sup) in microplate cultures. (**C** and **D**) Biofilm formation (**C**) and bacterial growth (**D**) of *E. faecalis* OG1RF in AUM (AUM), AUM supplemented with MRSA 1369 AUM culture supernatant (+MRSASup), and AUM supplemented with *K. pneumoniae* TOP52 AUM culture supernatant (+TOP52Sup) in microplate cultures. Data represent the mean ± SD derived from at least three independent experiments. Statistics were performed using a two-sided unpaired t test with $P \leq 0.05$ considered as statistically significant. $^*P \leq 0.05$, $^{**}P < 0.01$, $^{***}P < 0.001$, $^{****}P < 0.0001$, ns indicates not significant.

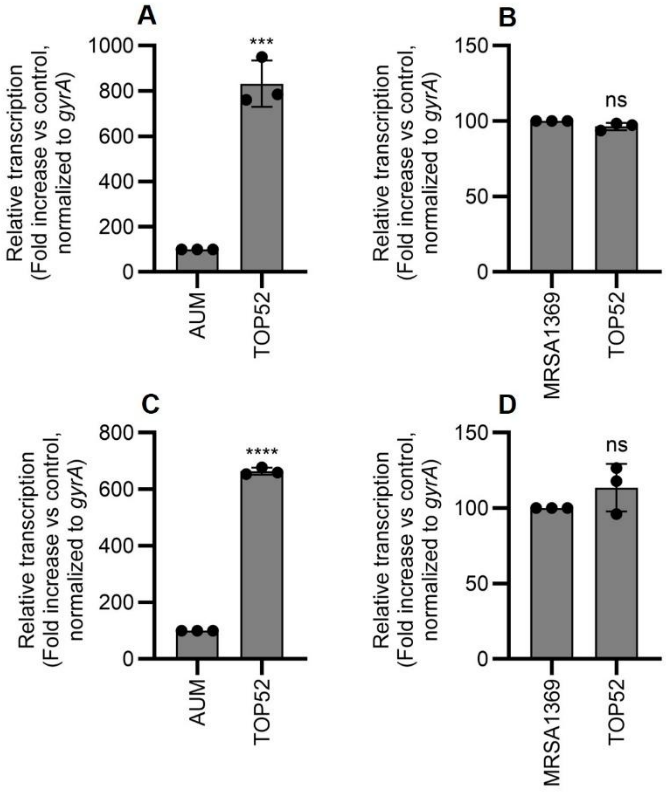


**Fig. S11. Expression of *gelE* and *sprE* genes in *E. faecalis* are affected by *K. pneumoniae*-produced active factors.** (**A** and **B**) *E. faecalis gelE* gene was up-regulated by *K. pneumoniae*-produced active factors at 9 hours (**A**) but not at 48 hours (**B**) post-inoculation as determined by RT-qPCR test. (**C** and **D**) *E. faecalis gelE* gene was up-regulated by *K. pneumoniae*-produced active factors at both 9 hours (**C**) and 48 hours (**D**) post-inoculation as determined by RT-qPCR test. Data represent the mean ± SD derived from three independent experiments. Statistics were performed using a two-sided unpaired t test with $P \leq 0.05$ considered as statistically significant. *$P \leq 0.05$, **$P < 0.01$, ***$P < 0.001$, ****$P < 0.0001$, ns indicates not significant.

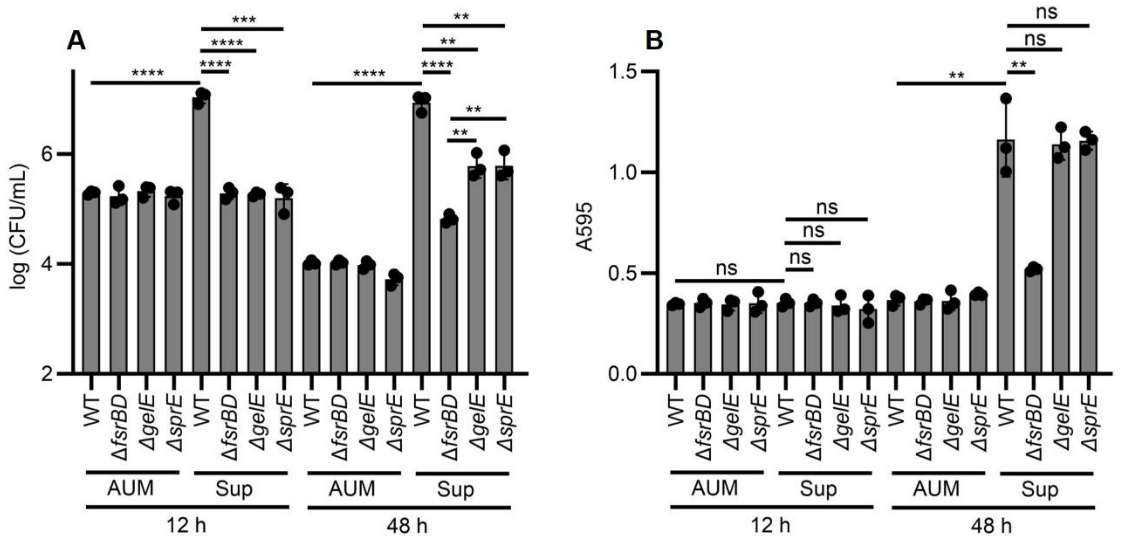


**Fig. S12. Bacterial growth and biofilm formation of wildtype *E. faecalis* and its isogenic mutants when supplemented with *K. pneumoniae*-produced factors.** (**A**) Bacterial growth of *E. faecalis* OG1RF (WT) and isogenic mutants Δ*fsrBD*, Δ*gelE*, and Δ*sprE* in AUM (AUM) and AUM supplemented with *K. pneumoniae* TOP52 supernatant (Sup) at 12 hours and 48 hours post-inoculation. (**B**) Biofilm formation of *E. faecalis* OG1RF (WT) and isogenic mutants Δ*fsrBD*, Δ*gelE*, and Δ*sprE* in AUM (AUM) and AUM supplemented with *K. pneumoniae* TOP52 supernatant (Sup) at 12 hours and 48 hours post-inoculation. Data represent the mean ± SD derived from three independent experiments. Statistics were performed using a two-sided unpaired t test with $P \le 0.05$ considered as statistically significant. *$P \le 0.05$, **$P < 0.01$, ***$P < 0.001$, ****$P < 0.0001$, ns indicates not significant.

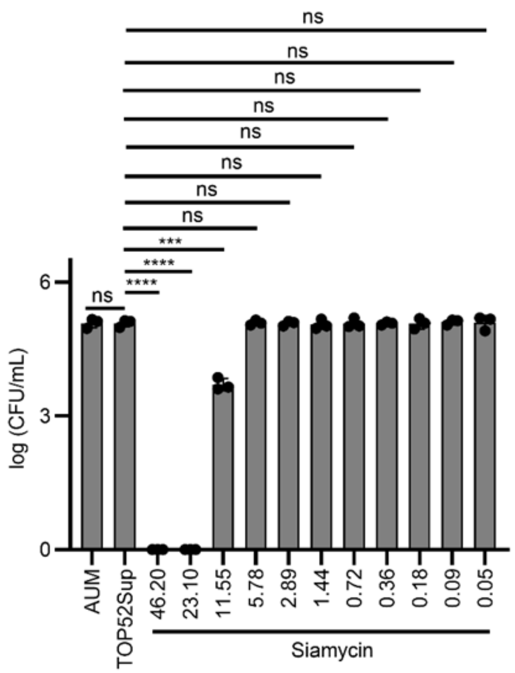


**Fig. S13.** Subinhibitory concentration of siamycin I against *E. faecalis* OG1RF cultured in AUM supplemented with *K. pneumoniae* TOP52 supernatant was determined by measuring bacterial growth (CFU/mL) at 3 hours post-inoculation, which identified 5.780 μM as the subinhibitory concentration. Data represent the mean ± SD derived from three independent experiments. Statistics were performed using a two-sided unpaired t test with $P \leq 0.05$ considered as statistically significant. *$P \leq 0.05$, **$P < 0.01$, ***$P < 0.001$, ****$P < 0.0001$, ns indicates not significant.

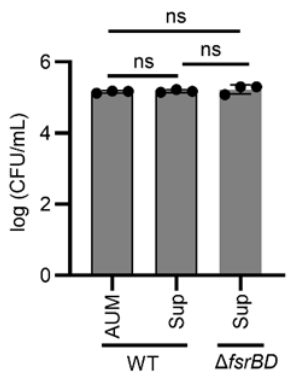


**Fig. S14.** Bacterial growth of *E. faecalis* OG1RF in AUM (WT-AUM) and AUM supplemented with *K. pneumoniae* TOP52 supernatant (WT-Sup), and *E. faecalis* OG1RFΔ*fsrBD* in AUM supplemented with *K. pneumoniae* TOP52 supernatant (Δ*fsrBD*-Sup), at 3 hours post-inoculation. Data represent the mean ± SD derived from three independent experiments. Statistics were performed using a two-sided unpaired t test with P ≤ 0.05 considered as statistically significant. *P ≤ 0.05, **P < 0.01, ***P < 0.001, ****P < 0.0001, ns indicates not significant.

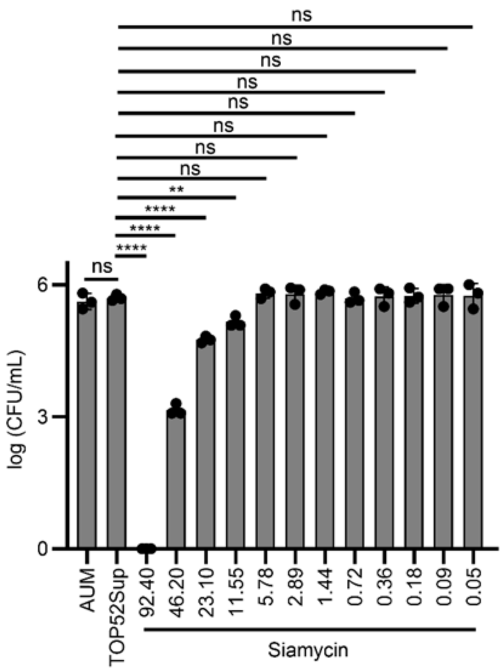


**Fig. S15.** Subinhibitory concentration of siamycin I against *E. faecalis* OG1RF cultured in AUM supplemented with *K. pneumoniae* TOP52 supernatant was determined by measuring bacterial growth (CFU/mL) at 9 hours post-inoculation, which identified 5.780 µM as the subinhibitory concentration. Data represent the mean ± SD derived from three independent experiments. Statistics were performed using a two-sided unpaired t test with $P \leq 0.05$ considered as statistically significant. *$P \leq 0.05$, **$P < 0.01$, ***$P < 0.001$, ****$P < 0.0001$, ns indicates not significant.

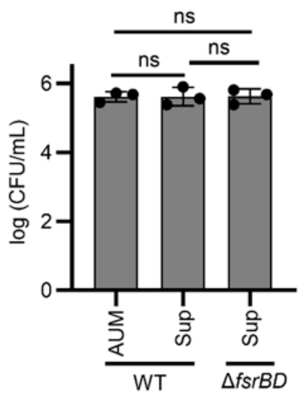


**Fig. S16.** Bacterial growth of *E. faecalis* OG1RF in AUM (WT-AUM) and AUM supplemented with *K. pneumoniae* TOP52 supernatant (WT-Sup), and *E. faecalis* OG1RFΔ*fsrBD* in AUM supplemented with *K. pneumoniae* TOP52 supernatant (Δ*fsrBD*-Sup), at 9 hours post-inoculation. Data represent the mean ± SD derived from three independent experiments. Statistics were performed using a two-sided unpaired t test with P ≤ 0.05 considered as statistically significant. *P ≤ 0.05, **P < 0.01, ***P < 0.001, ****P < 0.0001, ns indicates not significant.

### SUPPLEMENTARY TABLES

**Table S1. Average nucleotide identity (ANI) calculation for the 11 clinical *E. faecalis* isolates from Patient 85.**

| Patient 85_EF | | Orthologous Average Nucleotide Identity (OrthoANI) (%)[a] |
|---|---|---|
| Reference strain | Comparison strain | |
| EF00 | EF01 | 99.97 |
| | EF02 | 99.98 |
| | EF03 | 99.95 |
| | EF04 | 99.97 |
| | EF05 | 99.99 |
| | EF07 | 99.98 |
| | EF08 | 99.93 |
| | EF09 | 99.96 |
| | EF10 | 99.98 |
| | EF11 | 99.94 |

[a]ANI ≥ 95% indicates same species, and ANI ≥ 99.5% indicates same or nearly identical strain (*1, 2*).

**Table S2. Average nucleotide identity (ANI) calculation for the 11 clinical *K. pneumoniae* isolates from Patient 85.**

| Patient 85_KP | | Orthologous Average Nucleotide Identity (OrthoANI) (%)[a] |
|---|---|---|
| Reference strain | Comparison strain | |
| KP00 | KP01 | 99.99 |
| | KP02 | 99.99 |
| | KP03 | 99.99 |
| | KP04 | 99.99 |
| | KP05 | 99.99 |
| | KP06 | 99.99 |
| | KP07 | 99.98 |
| | KP08 | 99.98 |
| | KP09 | 99.99 |
| | KP10 | 99.99 |

[a]ANI ≥ 95% indicates same species, and ANI ≥ 99.5% indicates same or nearly identical strain (*1, 2*).

**Table S3. Average nucleotide identity (ANI) calculation for one phylogenetic group of clinical *E. faecalis* isolates from Patient 82 that includes 6 isolates.**

| Patient 82_EF-1 | | Orthologous Average Nucleotide Identity (OrthoANI) (%)[a] |
|---|---|---|
| Reference strain | Comparison strain | |
| EF01 | EF02 | 99.99 |
| | EF06 | 99.98 |
| | EF07 | 99.97 |
| | EF10 | 99.98 |
| | EF11 | 99.97 |

[a]ANI ≥ 95% indicates same species, and ANI ≥ 99.5% indicates same or nearly identical strain (*1, 2*).

**Table S4. Average nucleotide identity (ANI) calculation for one phylogenetic group of clinical *E. faecalis* isolates from Patient 82 that includes 3 isolates.**

| Patient 82_EF-2 | | Orthologous Average Nucleotide Identity (OrthoANI) (%) |
|---|---|---|
| **Reference strain** | **Comparison strain** | |
| EF03 | EF04 | 99.99 |
| | EF05 | 99.99 |

[a]ANI ≥ 95% indicates same species, and ANI ≥ 99.5% indicates same or nearly identical strain (*1*, *2*).

**Table S5. Average nucleotide identity (ANI) calculation for the 9 clinical *K. pneumoniae* isolates from Patient 82.**

| Patient 82_KP | | Orthologous Average Nucleotide Identity (OrthoANI) (%)[a] |
|---|---|---|
| Reference strain | Comparison strain | |
| KP02 | KP03 | 99.99 |
| | KP04 | 99.99 |
| | KP05 | 99.99 |
| | KP07 | 99.98 |
| | KP08 | 99.99 |
| | KP09 | 99.99 |
| | KP10 | 99.99 |
| | KP11 | 99.99 |

[a]ANI ≥ 95% indicates same species, and ANI ≥ 99.5% indicates same or nearly identical strain (*1, 2*).

**Table S6. Antibiotic resistance profile of 11 clinical *E. faecalis* isolates from Patient 85.**

| Isolate | Antibiotic resistance |
|---|---|
| 85EF00, 01, 02, 03 (N=55) | amikacin, ceftriaxone, chlortetracycline, ciprofloxacin, clindamycin, dalfopristin, demeclocycline, dibekacin, erythromycin, doxycycline, gentamicin, kanamycin, minocycline, netilmicin, oxytetracycline, pleuromutilin, quinupristin, rifampin, sisomicin, tetracycline, tobramycin, trimethoprim, vancomycin, 2'-N-ethylnetilmicin, 5-episisomicin, azithromycin, butirosin, larithromycin, dirithromycin, isepamicin, griseoviridin, lincomycin, lividomycin, madumycin II, neomycin, ostreogrycin B3, oleandomycin, paromomycin, patricin A, patricin B, pristinamycin IA, pristinamycin IB, pristinamycin IC, ribostamycin, roxithromycin, spiramycin, streptothricin, telithromycin, tylosin, vernamycin C, virginiamycin M1, virginiamycin S2 |
| 85EF04, 05, 07, 08, 09, 10, 11 (N=28) | amikacin, ceftriaxone, chlortetracycline, ciprofloxacin, clindamycin, dalfopristin, demeclocycline, dibekacin, erythromycin, doxycycline, gentamicin, kanamycin, minocycline, netilmicin, oxytetracycline, pleuromutilin, quinupristin, rifampin, sisomicin, tetracycline, tobramycin, trimethoprim, vancomycin, 2'-N-ethylnetilmicin, 5-episisomicin |

**Table S7. Antibiotic resistance profile of 11 clinical *K. pneumoniae* isolates from Patient 85.**

| Isolate | Antibiotic resistance |
| --- | --- |
| 85KP00, 01, 02, 03, 04, 05, 06, 07, 08, 09, 10 (N=48) | acriflavine, amikacin, ampicillin, azithromycin, benzalkonium chloride, cefaclor, cefalotin, cefdinir, cefditoren, cefepime, cefotaxime, cefoxitin, ceftazidime, ceftriaxone, chloramphenicol, chlorhexidine, ciprofloxacin, cloxacillin, colistin A, colistin B, defensin, ertapenem, erythromycin, fosfomycin, gentamicin, imipenem, kanamycin, metronidazole, nalidixic acid, neomycin, nitrofurantoin, norfloxacin, novobiocin, oxacillin, piperacillin, polymyxin B, puromycin, rifampin, spectinomycin, streptomycin, tetracycline, ticarcillin, tigecycline, tobramycin, triclosan, trimethoprim, vancomycin |

**Table S8. Antibiotic resistance profile of 9 clinical *E. faecalis* isolates from Patient 82.**

| Isolate | Antibiotic resistance |
|---|---|
| 82EF01, 02. 06, 07, 10, 11 (N=11) | acriflavine, ceftriaxone, ciprofloxacin, clindamycin, dalfopristin, erythromycin, pleuromutilin, quinupristin, rifampin, trimethoprim, vancomycin |
| 82EF03, 04, 05 (N=17) | acriflavine, ceftriaxone, ciprofloxacin, clindamycin, dalfopristin, erythromycinpleuromutilin, quinupristin, rifampin, trimethoprim, vancomycin, chlortetracycline, demeclocycline, doxycycline, minocycline, oxytetracycline, tetracycline |

**Table S9. Antibiotic resistance profile of 9 clinical *K. pneumoniae* isolates from Patient 82.**

| Isolate | Antibiotic resistance |
|---|---|
| 82EF02, 03, 04, 05, 07, 08, 09, 10, 11 (N = 48) | acriflavine; amikacin; ampicillin, azithromycin, benzalkonium chloride, cefaclor, cefalotin, cefdinir, cefditoren, cefepime, cefotaxime, cefoxitin, ceftazidime, ceftriaxone, chloramphenicol, chlorhexidine, ciprofloxacin, cloxacillin, colistin A, colistin B, defensin, ertapenem, erythromycin, fosfomycin, gentamicin, imipenem, kanamycin, metronidazole, nalidixic acid, neomycin, nitrofurantoin, norfloxacin, novobiocin, oxacillin, piperacillin, polymyxin B, puromycin, rifampin, spectinomycin, streptomycin, tetracycline, ticarcillin, tigecycline, tobramycin, triclosan, trimethoprim, vancomycin |

**Table S10. The list of most down- and up-regulated *E. faecalis* OG1RF genes with bacterial growth association induced by *K. pneumoniae* TOP52-produced functional factors.**

| Differentially expressed levels (DELs) | Differentially expressed genes (DEGs) |
|---|---|
| Most up-regulated | *agrBD, dnaC_1, gno_2, kduI1_2, mntB, mntH_2, pemK, pezT, yydJ, znuC_2, NA, NA, NA, NA, NA, NA, NA, NA, NA, NA, , NA, NA, NA, NA, NA, NA, NA, NA, NA, NA, NA, NA, NA, NA, NA, NA, NA, NA, NA, NA* |
| Most down-regulated | *fmnP, gspA_1, hmpT, ldh, mnmE, mnmG, nox, pdxK_1, pdxK_2, pflB, NA, NA, NA, NA* |

**Table S11. The list of most down- and up-regulated *E. faecalis* OG1RF genes with biofilm association induced by *K. pneumoniae* TOP52-produced functional factors.**

| Differentially expressed levels (DELs) | Differentially expressed genes (DEGs) |
|---|---|
| Most up-regulated | *agrBD, efp, exuR, glnQ_2, lacA_1, malG, murG, ntpC, phoP_1, pstS1_1, sigVL, stp_2, tetR_1, ytcD, NA, NA, NA, NA, NA, NA, NA, NA, NA, NA, NA, NA, NA* |
| Most down-regulated | *dnaC_1, hpf, hpt, infA, lrgA, lrgB, lytA_1, prgT_2, rpmE2, rpsL, ssrA, NA, NA, NA, NA, NA, NA, NA, NA, NA, NA, NA, NA, NA, NA, NA, NA, NA, NA, NA, NA, NA, NA* |

**Table S12. The transcription of gelE and sprE genes in *E. faecalis* OG1RF under the effect of *K. pneumoniae* TOP52-produced active factors.**

| Condition | Time (h) | Gene | -log(P)[c] | log(FC)[c] |
|---|---|---|---|---|
| Growth-enhancing | 9 h (AUM vs TOP52)[a] | *gelE* (KEFBALEM_02580) | 0.85 | -0.31 |
| | | *sprE* (KEFBALEM_02579) | 1.72 | -1.47 |
| Biofilm-enhancing | 48h (MRSA1369 vs TOP52)[b] | *gelE* (KEFBALEM_02580) | 0.72 | -0.41 |
| | | *sprE* (KEFBALEM_02579) | 4.21 | 1.41 |

[a]9 h (AUM vs TOP52): RNA-Seq analysis on *E. faecalis* OG1RF transcription grown in AUM only (AUM) and in AUM supplemented with *K. pneumoniae* TOP52 supernatant (TOP52) at 9 hours post inoculation.

[b]48 h (MRSA1369 vs TOP52): RNA-Seq analysis on *E. faecalis* OG1RF transcription grown in AUM supplemented with MRSA1369 supernatant (MRSA1369) and in AUM supplemented with *K. pneumoniae* TOP52 supernatant (TOP52) at 48 hours post inoculation.

[b]48 h (MRSA1369 vs TOP52): RNA-Seq analysis on *E. faecalis* OG1RF transcription grown in AUM supplemented with MRSA1369 supernatant (MRSA1369) and in AUM supplemented with *K. pneumoniae* TOP52 supernatant (TOP52) at 48 hours post inoculation.

[c]The criteria for determining differentially expressed genes are $|\log_2(FC)| > 2.0$ and $P < 0.05$ ($-\log(P) > 1.3$).

**Table S13. Strains used in this study.**

| Strain | Characteristic | Reference |
|---|---|---|
| 85EF00, 01, 02, 03, 04, 05, 07, 08, 09, 10, 11 | Eleven clinical *E. faecalis* (EF) isolates from Patient 85 with long-term implanted catheter | (*3*) |
| 85KP00, 01, 02, 03, 04, 05, 06, 07, 08, 09, 10 | Eleven clinical *K. pneumoniae* (KP) isolates from Patient 85 with long-term implanted catheter | (*3*) |
| 82EF01, 02, 03, 04, 05, 06, 07, 10, 11 | Nine clinical *E. faecalis* (EF) isolates from Patient 82 with long-term implanted catheter | (*3*) |
| 82KP02, 03, 04, 05, 07, 08, 09, 10, 11 | Nine clinical *K. pneumoniae* (KP) isolates from Patient 82 with long-term implanted catheter | (*3*) |
| *E. faecalis* OG1RF | Prototypical wildtype *E. faecalis* strain | (*4*) |
| *E. faecalis* OG1RFΔ*fsrBD* | *E. faecalis* OG1RF with in-frame deletion of gene *fsrBD* | This study |
| *E. faecalis* OG1RFΔ*gelE* | *E. faecalis* OG1RF with in-frame deletion of gene *gelE* | (*5*) |
| *E. faecalis* OG1RFΔ*sprE* | *E. faecalis* OG1RF with in-frame deletion of gene *sprE* | (*5*) |
| *K. pneumoniae* TOP52 | Prototypical wildtype *K. pneumoniae* strain | (*6*) |
| MRSA 1369 | Prototypical wildtype methicillin-resistant *S. aureus* strain | (*7*) |
